# Omicron-Enhanced Immunosuppressive Effects of SARS-CoV-2 ORF3a and ORF9b Accessory Proteins on Monocytic Inflammatory Response

**DOI:** 10.64898/2026.03.31.715496

**Authors:** Juozas Grigas, Miguel Padilla-Blanco, Unai Merino Herran, Blanca D López-Ayllón, Ana de Lucas Rius, Laura Mendoza Garcia, Raúl Fernández Rodríguez, Tránsito García García, Juan J Garrido, Arnoldas Pautienius, Arunas Stankevicius, Indre Kucinskaite-Kodze, Maria Montoya

## Abstract

This study investigates the poorly understood roles of SARS-CoV-2 accessory proteins using monocytic THP-1 cells expressing individual viral ORFs. ORF3a, ORF7b, and ORF9b were identified as major immunomodulators that suppress host inflammatory signaling. Specifically, cells expressing ORF3a or ORF9b exhibited reduced Toll-like receptor 4 (TLR4)-mediated production of key proinflammatory molecules—CCL2, CCL4, and IL-1β—resulting in diminished immune cell recruitment. Importantly, Omicron-associated mutations in ORF3a (T223I) and ORF9b (P10S⍰+⍰ΔE27N28A29) amplified this immunosuppressive effect, leading to stronger transcriptomic suppression consistent with Omicron’s reduced pathogenicity and clinical outcomes. These findings suggest that SARS-CoV-2 accessory proteins, particularly ORF3a and ORF9b, play pivotal roles in modulating monocytic immune responses. Enhanced suppression in Omicron variants highlights an evolutionary adaptation contributing to immune evasion and milder disease manifestations.

## INTRODUCTION

SARS-CoV-2 – the causative agent of coronavirus disease-19 (COVID-19) – is responsible for development of severe pneumonia, occasionally resulting in hypoxia and acute respiratory distress syndrome (1). Following severe COVID-19, many patients experience post-acute sequelae, known as ‘Long COVID’, resulting in a variety of symptoms affecting multiple organ systems (2). A combination of evasion of innate immune defenses (e.g. interferon response)(3), dysregulated immune and hyperinflammatory responses (4) have been proposed to contribute to the development of severe COVID-19. However, host innate immune response to SARS-CoV-2 differs depending on the predominant viral variant as well, with significantly lower plasma cytokine and chemokine levels reported in individuals infected with Omicron variant compared to those infected with either Wuhan strain or Delta variant (5), corresponding to lower severity of symptoms associated with Omicron infections (6).

Hyperinflammatory responses associated with severe COVID-19 may be linked via dysregulated inflammasome signaling pathways (1). While potential roles of structural SARS-CoV-2 proteins (spike, envelope, membrane and nucleocapsid) in inflammasome activation have previously been proposed, research on roles of other proteins in pathobiology of COVID-19, including accessory proteins (APs), has been limited. SARS-CoV-2 APs constitute eleven open reading frame (ORF) proteins: ORF3a, ORF3b, ORF3c, ORF3d, ORF6, ORF7a, ORF7b, ORF8, ORF9b, ORF9c and ORF10 (7). ORF3a has previously been described as a viroporin (8), capable of priming NLRP3 inflammasome resulting in proinflammatory cytokine and chemokine release via NF-κB pathway (9), reticulopathy (10,11) and autophagy (11,12) in infected cells. Multiple studies have previously indicated ORF3a association with inflammasome activation. However, its role in immune evasion – particularly related to type I interferon signaling pathway disruptions – remains controversial (9,13). While ORF9b association with interferon antagonism and NLRP3 activation has been well established (14–17), capacity of ORF9b to suppress expression of multiple pro-inflammatory mediators, including IL6, TNF, CCL2 and CXCL10, has also been proposed (14,18). One study suggests that ORF9b may be a modulating rather than a strictly inhibiting protein, up- or downregulating host immune response pathways to favor viral retention (19). Similarly, ORF7b has previously been described as capable of interferon antagonism (20–22), while investigations into its other roles related to host immune responses, including inflammatory response, have largely been inconclusive (21,23,24).

SARS-CoV-2 infections by currently dominant Omicron strains are characterized by milder symptoms and decreased pathogenicity (25). Similarly, interactions between viral and host proteins differ depending on the variant of origin, with APs of Omicron strains demonstrating reduced levels of alterations typically associated with SARS-CoV-2 pathobiology (26). While most significant mutations between SARS-CoV-2 strains have been observed in the Spike protein, different Omicron variants are characterized by a variety of unique mutations at AP-coding genes, including ORF3a, ORF6, ORF7a, ORF7b, ORF8 and, most notably, ORF9b (27). Mutations exhibited by ORF3a of Omicron variants (e.g. I35T and Y107H) have been proposed as contributing to reduced ORF3a-dependant immune response activation in T cells (28), while mutation such as T223I has been associated with increased effect on NF-κB-mediated cytokine production (29). Despite the data suggesting increased expression of ORF9b in Omicron-infected cells (30), very few studies investigated changes related to Omicron mutations in host immune response, finding little difference between Omicron and Wuhan variants (31).

While multiple studies investigating the effect of SARS-CoV-2 APs in epithelial cells are available, including lung and kidney cells, the number of studies investigating their impact on monocytes/macrophages, being the primary effector cells involved in COVID-19 pathobiology, is limited. Severe SARS-CoV-2 cases are accompanied by excessive amounts of pro-inflammatory cytokines (e.g. IL-1α, IL-1β, TNF) and chemokines (e.g. CCL2, CXCL10) released by monocytes/macrophages (32). Exaggerated inflammation resulting from ACE2 downregulation has been well established. However, recent data suggests an additional trigger associated with SARS-CoV-2 Spike and TRL4 interaction (33). Persistent or abnormal activation of proinflammatory pathways in monocytes/macrophages has also been proposed as the major contributing factor to the development Long COVID (32). NLRP3 inflammasome activation, characterized by ASC speck assembly, has been visualized in ORF3a-transduced human macrophage-like THP-1-ASC-GFP reporter cells (34). Additionally, increased expression of TNF-α, IL-1β, IL-6 and IL-8 in ORF3a expressing monocytic THP-1 cells (9,35) has previously been reported, suggesting ORF3a role in NLRP3 inflammasome activation in monocytes/macrophages. Interestingly, while ORF9b has been associated with interferon antagonism and suppressed expression of multiple pro-inflammatory mediators in epithelial cells, increased activity of caspase-1 – expression of which follows NLRP3 inflammasome activation and leads to pro-inflammatory cytokine release – has been reported in THP-1 cells transiently transduced with ORF9b and activated with LPS and ATP compared to their wild type counterparts (17). While THP-1 cell transduction with ORF7b has also been carried out before, significant ORF7b-associated alterations to immune response pathways have not been detected (17). Therefore, while limited data investigating SARS-CoV-2 APs impact on monocyte/macrophage cells largely reveal similarities with epithelial cells – especially with regards to ORF3a – not enough information is currently available to draw definitive conclusions.

The main objective of this study was to elucidate the impact of SARS-CoV-2 APs from Wuhan and Omicron variants on the immune response in human monocyte/macrophage cells. Individual SARS-CoV-2 ORF3a, ORF6, ORF7a, ORF7b, ORF8, ORF9b, ORF9c and ORF10-expressing human lung adenocarcinoma (A549-ORF) and monocytic (mTHP-1-ORF) cells have been constructed and gene expression changes in each cell type have been identified. mTHP-1-ORFs exhibiting the most significant alterations in immune response-related pathways were selected for further analysis. APs derived from both Wuhan (Wuhan-Hu-1) and Omicron (21L (BA.2)) lineages were analyzed where appropriate in monocytic and macrophage-like THP-1 cells (MTHP-1-ORF). The production, secretion and functional impact of the most significantly altered genes and proteins were subsequently assessed. Finally, our findings from *in vitro* cell model were compared with transcriptomics data from bronchoalveolar lavage (BAL) samples of COVID-19 patients to explore potential links between SARS-CoV-2 AP roles and immune response alterations.

## RESULTS

### A549 and mTHP-1 cells individually transduced with SARS-CoV-2 APs exhibit distinct transcriptomic profiles

Due to respiratory system cells being the primary targets of SARS-CoV-2, lung carcinoma A549 and similar cell lines have previously been used for studying SARS-CoV-2 APs, demonstrating upregulation of immune response-associated pathways, such as inflammasome activation, associated with some APs (36,37). In order to compare and see if similar AP-associated responses can be detected in the primary effector cells involved in COVID-19 pathology, such as monocyte/macrophage cells, A549 and monocytic leukemia THP-1 cells (mTHP-1) were individually transduced with OR3a, ORF6, ORF7a, ORF7b, ORF8, ORF9b, ORF9c and ORF10 and selected for expression of viral genes as previously described (38)(Supplementary Fig. 1). For further detection, all APs were fused to a 2x Strep-tag (ST) in their N- or C-terminal domain and their expression was confirmed by RT-qPCR (Fig. 1A). Transcriptome analysis of AP-expressing cells was performed using mRNA sequencing.

**Fig. 1.**
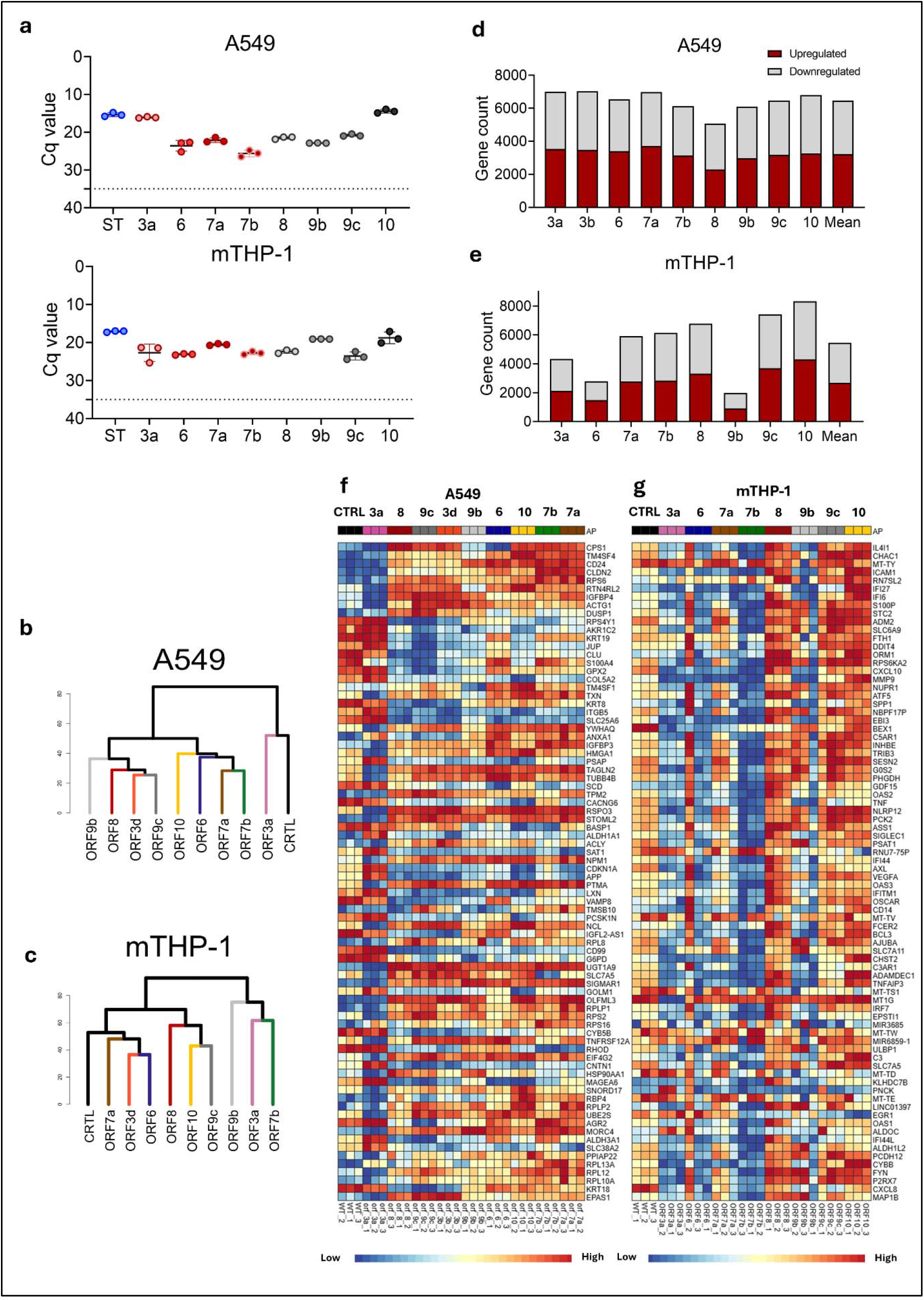
Transcriptome analysis of SARS-CoV-2 accessory protein-expressing A549 and THP-1 cells. **a**. SARS-CoV-2 AP mRNA expression in respective ORF-transduced A549 and mTHP-1 cells. ST refers to 2x Strep-tag confirmation in control cells, transduced with lentiviral vector only. Cq value threshold 35 was assumed based on negative control data. Error bars represent 95% CI values. **b**. Distance clustering dendrograms of A549-ORF and control cells, and **c**. mTHP-1-ORF and mTHP-1 control cells based on the expression levels of altered genes. Dendrograms constructed based on FPKM values additionally normalized with variance-stabilizing transformation and analyzed using DESeq2 package in R. **d**. Bar plots of A549-ORF3a vs A549 control group and **e**. mTHP-1-ORF3a vs mTHP-1 control group DEGs, demonstrating upregulation (red) or downregulation (grey). Only significantly altered (adjusted *p*-value < 0.05) genes were included. **f**. Heatmaps showing top 80 significant DEGs in A549-ORF (3a, 3b, 6, 7a, 7b, 8, 9b, 9c and 10) vs A549 controls (WT) and **g**. mTHP-1-ORF vs mTHP-1 controls. Data normalized using Log2-counts per million (logCPM) transformation, while differential expression carried out by DESeq2 package in R. The heatmap was generated by the visualization module in ExpressAnalyst.

The differential gene expression analysis was carried out by comparing A549-ORF and mTHP-1-ORF cells with their respective vector-transduced cells (controls-ST). Distance clustering analysis based on gene expression levels revealed two distinct clusters in A549 cells, one of which consisted of control and A549-ORF3a, while the other encompassed all other tested A549-ORFs (Fig. 1B). Clustering was markedly different in mTHP-1 cells, characterized by two clusters, one of which encompassed ORF9b-, ORF3a- and ORF7b-transduced cells (Fig. 1C). Full gene count and clustering data for A549-ORF and mTHP-1-ORF cells available in Supplementary Fig. 2 and 3, respectively.

Significant differentially expressed genes (DEGs) were selected for further analysis based on adjusted *p*-value (*q*-value < 0.05) and log2-fold change (|Log2FC| > 1.5). We have observed that on average, more genes were identified in A549-ORF vs A549 control groups (DEGs *n* = 6458) compared to mTHP-1-ORF vs mTHP-1 control groups (DEGs *n* = 5465) (Fig. 1D, 1E). Overall, both A549-ORF vs A549 control and mTHP-1-ORF vs mTHP-1 control differential expression was characterized by a similar ratio of upregulated to downregulated genes (average 49.75% upregulated vs 50.25% downregulated DEGs in A549-ORF vs A549 control groups (Fig. 1D), average 49.09% upregulated vs 50.91% downregulated DEGs in mTHP-1-ORF vs mTHP-1 control groups (Fig. 1E)). Distinct DE profiles were observed in A549 and THP-1 cells, with only two overlapping genes (RPS6, SLC7A5) within the 80 most significant DEGs, suggesting vastly different effects of SARS-CoV-2 APs in A549 (Fig. 1F) and mTHP-1 cells (Fig. 1G).

### Inflammatory response-related pathways were significantly altered by SARS-CoV-2 ORF3a, ORF7b and ORF9b in mTHP-1 cells

Since transcriptomic profiles of SARS-CoV-2 ORF-expressing mTHP-1 cells differed significantly from A549-ORFs, we further focused on mTHP-1-ORFs in order to identify APs associated with the most significant alterations. Pathway enrichment analysis (39) revealed ORF3a, ORF7b and ORF9b as the most significantly involved APs within the top 20 most affected pathways in mTHP-1-ORFs (Fig. 2A). In turn, we selected these three APs for subsequent analyses. A significantly higher number of DEGs (adjusted *p*-value < 0.05, |Log2FC| > 1.5) were present in mTHP-1-ORF3a and mTHP-1-ORF7b cells, compared to mTHP-1-ORF9b cells (508, 535 and 129, respectively; Fig. 2B). While a substantial amount of DEGs were exclusive for mTHP-1-ORF3a and mTHP-1-ORF7b cells (266 and 289, respectively), most DEGs in mTHP-1-ORF9b overlapped with either mTHP-1-ORF3a (*n*=18), mTHP-1-ORF7b (*n*=22), or both (*n*=50)(Fig. 2B).

**Fig. 2.**
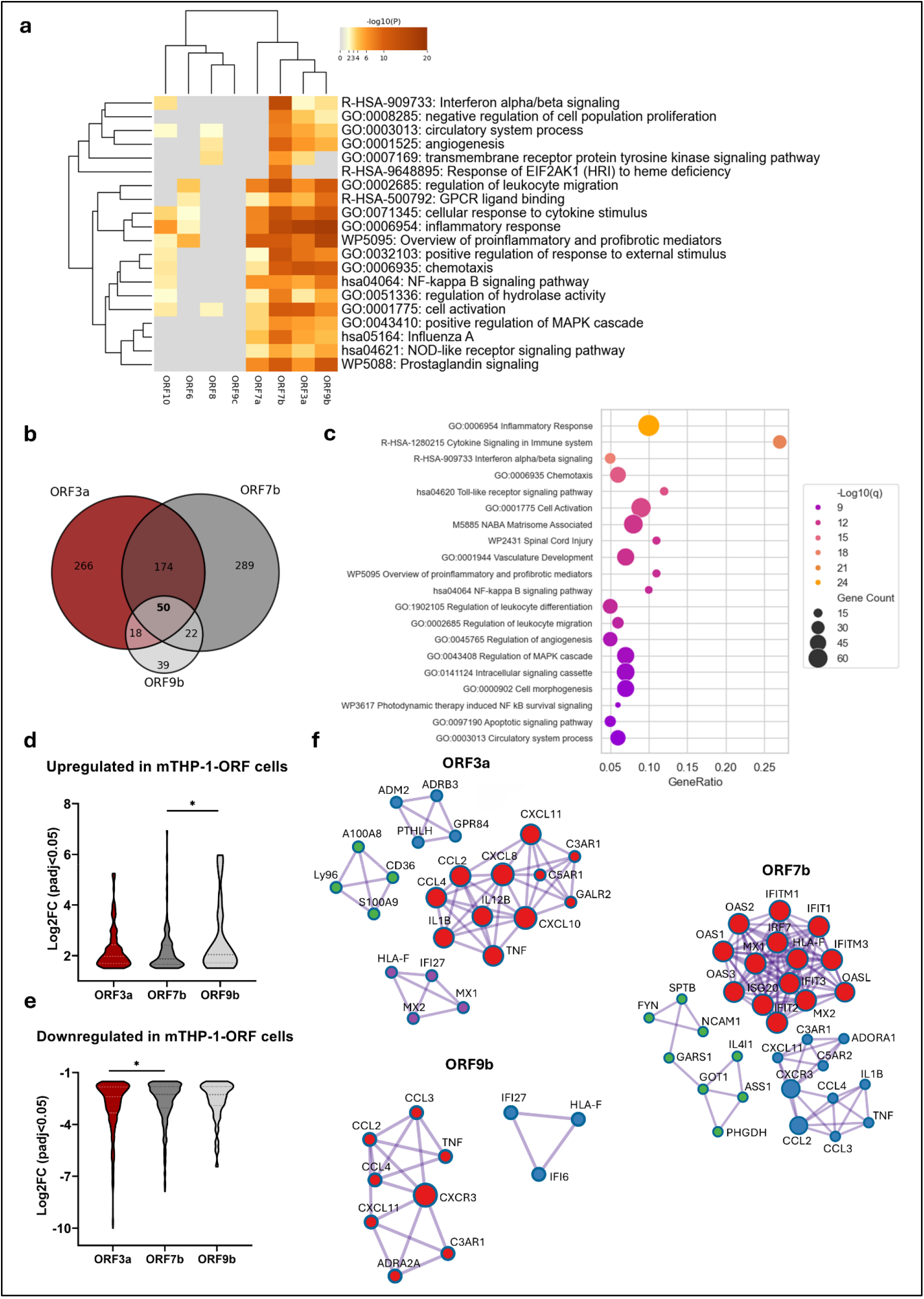
Pathway enrichment analysis of SARS-CoV-2 accessory protein-expressing THP-1 cells, based on KEGG Pathway, GO Biological Processes, Reactome Gene Sets, Canonical Pathways, CORUM, WikiPathways, and PANTHER Pathway ontology sources. **a**. Heatmap of top 20 enriched terms across significant DEGs (adjusted *p*-value < 0.05, |Log2FC| > 1.5) in mTHP-1-ORF cells. Analysis carried out using Metascape Gene List Analysis tool. **b**. Venn diagram demonstrating all significant DEGs (adjusted *p*-value < 0.05) between mTHP-1 cells transduced with SARS-CoV-2 ORF3a, ORF7b and ORF9b. **c**. Dot plot of pathway-enrichment analysis of significant DEGs (adjusted *p*-value < 0.05, |Log2FC| > 1.5) in mTHP-1-ORF3a, -ORF7b and -ORF9b cells, highlighting the gene count and ratio to all significant DEGs that were found in the given ontology term. **d**. Violin plots demonstrating log2-fold change values of all significantly upregulated (adjusted *p*-value < 0.05, Log2FC > 1.5) and **e**. downregulated (adjusted *p*-value < 0.05, Log2FC < -1.5) DEGs in mTHP-1-ORF3a, -ORF7b and -ORF9b cells. *p*-values were calculated using unpaired two-tailed t-test, \**p*<0.05. **f**. Protein-protein Interaction Enrichment Analysis with Molecular Complex Detection (MCODE) algorithm applied to identify densely connected network components in mTHP-1-ORF3a, -ORF7b and -ORF9b cells. Only physical interactions in STRING (physical score > 0.132) and BioGrid are used. Each node corresponds to a gene involved in a given MCODE cluster. Analysis carried out using Metascape Gene List Analysis tool.

Evaluation of most significantly affected pathways based on KEGG Pathway, GO Biological Processes, Reactome Gene Sets, Canonical Pathways, CORUM, WikiPathways, and PANTHER Pathway ontology sources in mTHP-1-ORF3a, -ORF7b and -ORF9b cells revealed an overrepresentation of pathways associated with immune responses, including *GO:0006954 Inflammatory Response, R-HSA-1280215 Cytokine Signaling in Immune system, R-HSA-909733 Interferon alpha/beta signaling, GO:0006935 chemotaxis, hsa04620 Toll-like receptor signaling pathway* and *hsa04064 NF-kappa B signaling pathway* (Fig. 2C). These results in turn suggest that ORF3a, ORF7b and ORF9b primarily interact with targets that alter immune response-related pathways in mTHP-1 cells.

Upon closer inspection of significant DEGs (adjusted *p*-value < 0.05) in mTHP-1-ORF3a, -ORF7b and - ORF9b cells, we observed both upregulated (Fig. 2D) and downregulated (Fig. 2E) genes. Similar distribution trends were observed between mTHP-1-ORF3a, -ORF7b and -ORF9b cells, with some genes demonstrating Log2FC of as low as -10 in mTHP-1-ORF3a, suggesting an inhibitory capacity of these ORFs (Fig. 2E). In order to identify putative intracellular targets of ORF3a, ORF7a and ORF9b in mTHP-1 cells, Protein-protein Interaction Enrichment Analysis with Molecular Complex Detection (MCODE) algorithm was applied to generate clusters of associated DEGs (Fig. 2F). Multiple clusters, primarily constituting genes involved in the immune response function of the cell, were identified.

Six significant DEGs (*q*-value < 0.05) forming MCODE clusters overlapped in mTHP-1-ORF3a, -ORF7b and ORF9b cells (*CCL2, CCL4, CXCL11, C3AR1, TNF* and *HLA-F;* Fig. 3A). Moreover, *IL1B, MX1* and *MX2* overlapped in THP-1-ORF7b and -ORF3a cells, while *CCL3* and *CXCR3* overlapped in THP-1-ORF9b and -ORF7b cells (Fig. 3A). Finally, *IFI27* overlapped in mTHP-1-ORF9b and -ORF3a cells (Fig. 3A). Interestingly, overlapping genes were downregulated in mTHP-1-ORF3a, ORF7b and ORF9b cells (Fig. 3B), further confirming inhibitory capacity of these ORFs, targeting factors associated with immune response-related pathways, in turn suggesting putative blunting of immune response in mTHP-1 cells by ORF3a, ORF7b and ORF9b APs.

**Fig. 3.**
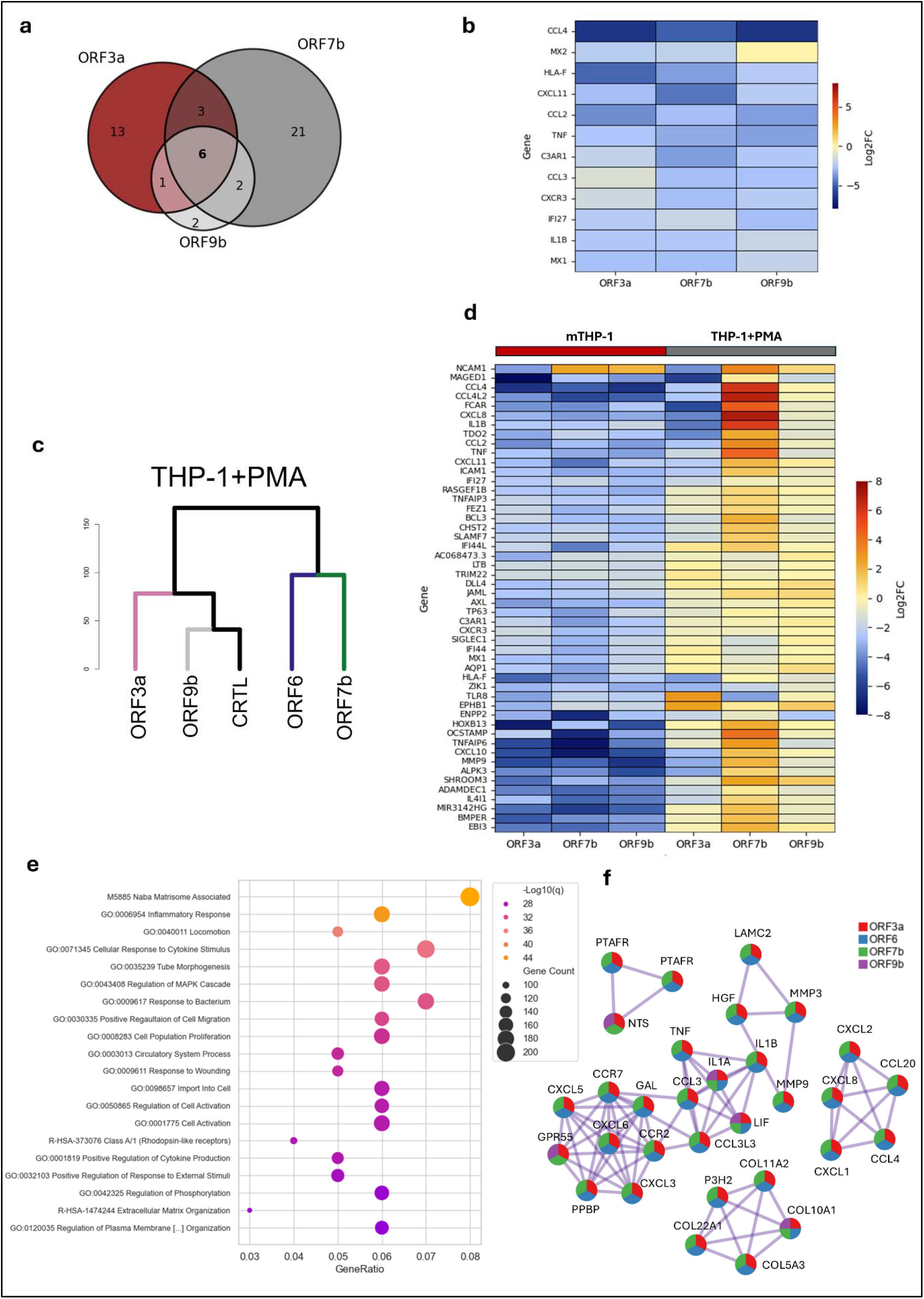
Immune response-related genes in SARS-CoV-2 ORF3a-, ORF7b- and ORF9b-transduced mTHP-1 cells and differences observed in MTHP-1 cell transcriptome. **a**. Venn diagram demonstrating all significant DEGs (adjusted *p*-value < 0.05) forming MCODE clusters between mTHP-1 cells transduced with SARS-CoV-2 ORF3a, ORF7b and ORF9b. **b**. Heatmap showing alterations in significant DEGs (adjusted *p*-value < 0.05) forming MCODE clusters in mTHP-1 cells transduced with SARS-CoV-2 ORF3a, ORF7b and ORF9b. Log2-fold change values were acquired using DESeq2 package in R. **c**. Distance clustering dendrogram of control MTHP-1 cells and SARS-CoV-2 ORF3a-, ORF7b-, ORF9b- and ORF6-transduced MTHP-1 cells based on the expression levels of altered genes. Dendrogram constructed based on FPKM values additionally normalized with variance-stabilizing transformation and analyzed using DESeq2 package in R. **d**. Heatmap showing alterations in 50 overlapping genes of mTHP-1-ORF3a, - ORF7b and -ORF9b cells (Fig. 2B) in ORF3a-, ORF7b- and ORF9b-transduced mTHP-1 and MTHP-1 cells. Log2-fold change values were acquired using DESeq2 package in R. **e**. Dot plot of pathway-enrichment analysis of significant DEGs (adjusted *p*-value < 0.05, |Log2FC| > 1.5) in MTHP-1-ORF3a, -ORF7b and - ORF9b cells, highlighting the number of genes with membership in the given ontology term and ratio of all significant DEGs that are found in the given ontology term. **f**. Protein-protein Interaction Enrichment Analysis MCODE algorithm applied to identify densely connected network components in MTHP-1-ORF3a, -ORF6, -ORF7b and -ORF9b cells. Only physical interactions in STRING (physical score > 0.132) and BioGrid are used. Each node corresponds to a gene involved in a given MCODE cluster. Analysis carried out using Metascape Gene List Analysis tool.

### Downregulated transcriptome profiles of mTHP-1 and MTHP-1 cells expressing ORF3a, ORF7b and ORF9b of Wuhan origin

Since monocytes/macrophages play an important role in SARS-CoV-2 pathobiology, with both monocytes and lung resident macrophages exhibiting inflammasome activation (40), we aimed to evaluate if the changes in mTHP-1 cells following transduction with SARS-CoV-2 ORF3a, ORF7b and ORF9b were observed in MTHP-1-ORFs as well. mTHP-1-ORF differentiation into macrophage-like cells was induced with phorbol 12-myristate-13-acetate (PMA). In addition to controls, mTHP-1 and MTHP-1 cell expressing ORF6 were used as an outgroup control. Distance clustering analysis based on gene expression levels revealed different clustering trends in MTHP-1-ORFs (Fig. 3C) compared to mTHP-1-ORFs (Fig. 1C). Clustering trends in MTHP-1 were somewhat similar to A549-ORFs (Fig. 1B) where A549-ORF7b clustered close to A549-ORF6, while A549-ORF3a-transduced cells formed a distinct cluster with controls. Similarly, ORF3a-, ORF9b-transduced and control MTHP-1 cells constituted a separate cluster from ORF7b- and ORF6-transduced MTHP-1 cells. Fifty overlapping DEGs in mTHP-1-ORF3a, -ORF7b and - ORF9b cells (Fig. 2B) were compared to their counterparts in MTHP-1-ORF3a, -ORF7b and -ORF9b, revealing significantly different profiles (Fig. 3D). While strong downregulation in immune system-related genes (e.g. *TNF, CCL4, IL1B, CXCL11* and *CCL2*) was observed in mTHP-1-ORF3a, -ORF7b and -ORF9b cells, MTHP-1-ORF3a and particularly MTHP-1-ORF9b demonstrated a more modest downregulation. Interestingly, significantly more upregulated genes were present in MTHP-1-ORF7b cells compared to their monocyte-like counterparts (Fig. 3D).

Although pathway enrichment analysis of MTHP-1-ORF3a, -ORF6, -ORF7b and -ORF9b cells revealed significant alterations in a greater variety of unrelated pathways (Fig. 3E), compared to respective mTHP-1-ORFs (Fig. 2C), a substantial number of most significantly altered pathways were immune response-related pathways, including *GO:0006954 Inflammatory Response, GO:0071345 Cellular Response to Cytokine Stimulus, GO:0009617 Response To Bacterium, GO:0030335 Positive Regulation of Cell Migration* and *GO:0001819 Positive Regulation of Cytokine Production* (Fig. 3E). MCODE analysis further demonstrated that clusters with the most substantial number of members were comprised of immune response-related genes, such as *IL1B, CCL3, CXCL5, TNF* or *CXCL8, CCL4, CXCL2* (Fig. 3F).

### Expression of ORF3a or ORF9b proteins derived from SARS-CoV-2 Omicron variants leads to downregulation of genes associated with the inflammatory response in both mTHP-1 and MTHP-1 cells

Given the observed downregulation of multiple immune response-associated genes in SARS-CoV-2 ORF-transduced mTHP-1-ORFs, we next examined whether mutations characteristic of ORFs present in most SARS-CoV-2 Omicron lineages exerted different effects on gene expression. Mutations in different Omicron lineages have previously been proposed to potentially contribute to lower pathogenesis compared to other variants of concern (41,42). To evaluate the effect of ORF mutations in Omicron lineages, mTHP-1 cells were transduced with ORF3a and ORF9b harboring the most conserved Omicron mutations, namely T223I and P10S+ΔE27N28A29, respectively (Fig. 4A, Supplementary Fig. 4). These mutations emerged in Omicron lineages at the beginning of 2022 and are still present in the currently circulating variants. ORF7b does not harbor any conserved mutations since early emergence of Omicron (7). Therefore, Omicron-specific ORF7b transduction was not possible. While genetically similar, mTHP-1 cells transduced with Wuhan or Omicron ORF3a shared alterations in 242 genes compared to the total of 508 and 499 significant DEGs in mTHP-1-ORF3a Wuhan and Omicron cells, respectively, while alterations in 79 genes were shared compared to the total of 1318 and 209 significant DEGs in MTHP-1-ORF3a Wuhan and Omicron cells, respectively (Fig. 4B). Similar trend was observed even more dramatically with respect to ORF9b, with shared alterations in only 66 genes compared to the total of 129 and 350 significant DEGs present in mTHP-1-ORF9b Wuhan and Omicron cells, respectively, and shared alterations in 24 genes compared to the total of 131 and 692 significant DEGs in MTHP-1-ORF9b Wuhan and Omicron cells, respectively (Fig. 4C).

**Fig. 4.**
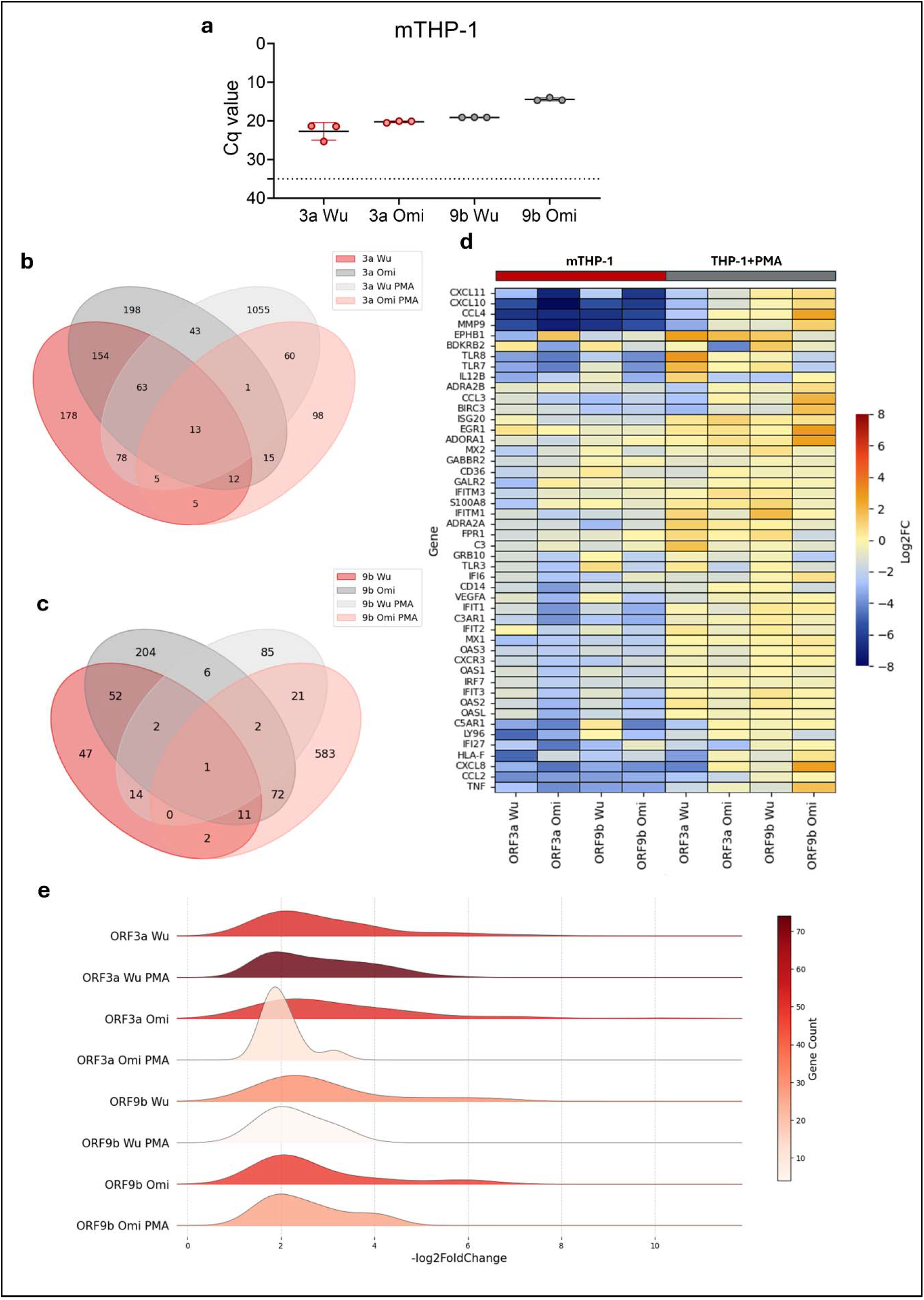
Transcriptome analysis of mTHP-1 cells transduced with SARS-CoV-2 Omicron 21L (BA.2) ORF3a and ORF9b. **a**. SARS-CoV-2 Wuhan and Omicron AP mRNA expression in respective ORF-transduced mTHP-1 cells. Cq value threshold 35 was assumed based on negative control data. Error bars represent 95% CI values. **b**. Venn diagrams demonstrating all significant DEGs (adjusted *p*-value < 0.05) between mTHP-1 and MTHP-1 cells transduced with Wuhan and Omicron ORF3a and **c**. ORF9b. **d**. Heatmap showing 48 genes previously found to form clusters according to MCODE analysis (Fig. 2F, Fig. 3F) in Wuhan and BA.2 ORF3a- and ORF9b-transduced mTHP-1 and MTHP-1 cells. Log2-fold change values were acquired using DESeq2 package in R. **e**. Ridgeline plot showing significantly downregulated (adjusted *p*-value < 0.05) genes involved in *GO:0006954 Inflammatory Response* pathway in Wuhan and BA.2 ORF3a- and ORF9b-transduced mTHP-1 and MTHP-1 cells.

Comparing alterations in mTHP-1 and MTHP-1 cells transduced with Wuhan and Omicron ORFs with respect to genes previously found to form clusters as per MCODE analysis (Fig. 2F, Fig. 3F), substantially greater downregulation has been observed in Omicron ORF-transduced mTHP-1 genes associated with inflammatory and interferon response pathways, such as *CXCL11, CXCL10, CCL4, TLR8, TLR7, TLR3, IFIT1, C3AR1, IFIT2, IFIT3, C5AR1, CCL2* and *TNF* (Fig. 4D), compared to Wuhan ORF-transduced mTHP-1 cells. However, this trend was largely absent in MTHP-1 cells, with some marked exceptions, such as downregulation of *BDKRB2, IFI27, CCL2, TLR3* and *IL12B* in MTHP-1-ORF3a Omicron cells and *TLR7, TLR8, CD14* and *FPR1* in MTHP-1-ORF9b Omicron cells (Fig. 4D).

In order to investigate the impact of Wuhan and Omicron ORF9b and ORF3a in mTHP-1 and MTHP-1 cells on all genes associated with inflammatory response, genes involved in *GO:0006954 Inflammatory Response* pathway were compared. Significantly downregulated genes were selected for each cell type. While in most MTHP-1-ORFs, a lower number of inflammatory response genes were downregulated compared to their monocytic counterparts, a significantly higher number of inflammatory response genes was observed in MTHP-1-ORF3a Wuhan cells compared to mTHP-1-ORF3a Wuhan cells (Fig. 4E). More downregulated inflammatory response genes in Omicron ORF9b-transduced mTHP-1 and MTHP-1 cells were observed, while that was true only for mTHP-1-ORF3a but not MTHP-1-ORF3a cells (Fig. 4E). Moreover, while none of the downregulated gene Log2FC values of Wuhan ORF-transduced MTHP-1 cells exceeded -4, some genes of both Omicron ORF3a- and ORF9b-transduced MTHP-1 cells did (Fig. 4E). Overall, while most downregulated inflammatory response genes in MTHP-1-ORFs clustered around Log2FC values of -2, distribution of these values was more heterogenous among mTHP-1-ORF cells, with some genes reaching values below -6.

### Expression of SARS-CoV-2 ORF3a and ORF9b in mTHP-1 cells modulated cytokine and chemokine production, leading to altered myeloid cell recruitment

Among the cellular mechanisms involved in the activation of the inflammatory response, TLR4-dependent signaling pathway plays a central role. Its activation ultimately promotes the nuclear translocation of NF-κB, leading to the production of proinflammatory cytokines and chemokines (43). Therefore, we stimulated the TLR4-dependent pathway with lipopolysaccharide (LPS) in our control and ORF3a-, ORF6-, ORF7b- and ORF9b-transduced mTHP-1 cells (with ORF3a and ORF9b of both Wuhan and Omicron origin) (Fig. 5A). Similarly to transcriptomic results which demonstrated *IL1B, CCL2, CCL4* and *CXCL11* reversal to baseline expression (Log2FC ≈ 0) in MTHP-1-ORF9b cells (Fig. 5B), LPS stimulation of mTHP-1-ORF9b cells resulted in a varying cytokine/chemokine mRNA expression. While *CCL2, CXCL11* and *IL1B* expression did not significantly differ from control cells at both 100 ng/ml and 400 ng/ml LPS stimulation, a significant decrease of *CCL4* expression was observed in cells expressing Omicron ORF3a (Fig. 5C). Combined data from both LPS concentrations revealed a significant difference of *CCL2, CCL4* and *IL1B* mRNA expression between Wuhan and Omicron ORF9b-transduced cells, demonstrating downregulation in Omicron ORF9b expressing mTHP-1 cells (Fig. 5C). These findings are in line with alterations observed in cytokine/chemokine production, where Wuhan ORF9b expressing cells primarily demonstrated a decrease in CCL2, CCL4 and CXCL11 that did not reach statistical significance, while Omicron ORF9b expressing mTHP-1 cells demonstrated significant decrease of CCL2, CCL4 and IL-1β production at 100 ng/ml LPS (Fig. 5D). As expected, both LPS-untreated control and mTHP-1-ORF9b cells exhibited marginal proinflammatory cytokine/chemokine mRNA expression and production (Supplementary Fig. 5A). In line with cytokine mRNA expression and production results, mTHP-1 cell movement through a Transwell membrane was significantly decreased in the presence of both LPS-treated Wuhan and Omicron mTHP-1-ORF9b growth media (Fig. 5E). CCL2 contributed to the majority of mTHP-1 cell mobility, resulting in a significant CCL2-associated movement decrease in the presence of LPS-treated Wuhan mTHP-1-ORF9b growth media, and further decrease in Omicron mTHP-1-ORF9b growth media (Fig. 5F). In combination, this data suggests a modest downregulation in Wuhan ORF9b-transduced mTHP-1 cells in the presence of TLR4 stimulation, with a significantly stronger downregulation in Omicron ORF9b-transduced mTHP-1 cells in the presence of TLR4 stimulation, resulting in a suppressed cytokine/chemokine production.

**Fig. 5.**
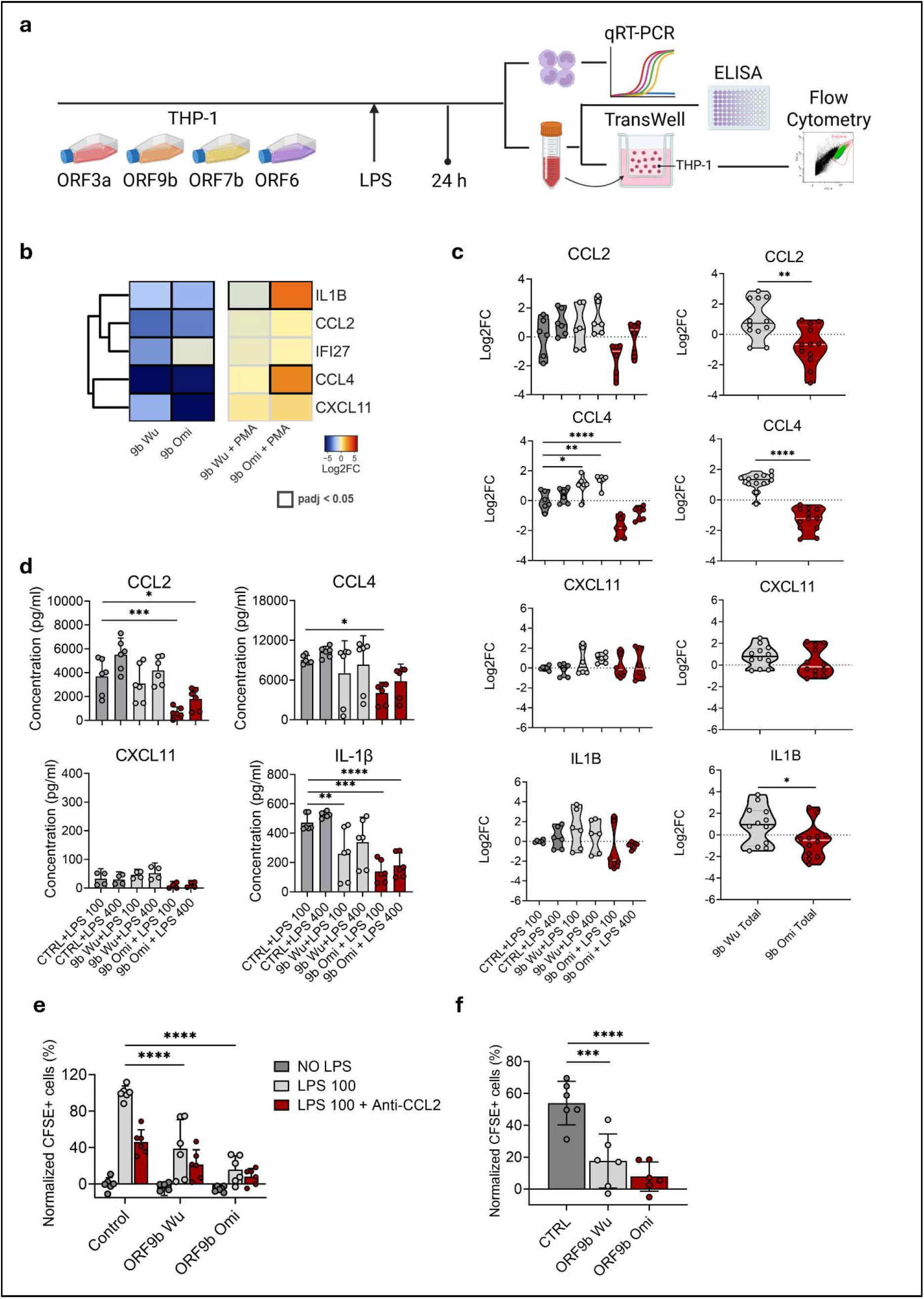
Proinflammatory cytokine/chemokine expression and production changes in ORF9b-expressing mTHP-1 cells following TLR4 stimulation. **a**. Schematic of the experimental pipeline. SARS-CoV-2 ORF expressing mTHP-1 cells were stimulated with 100 ng/ml, 400 ng/ml of LPS or mock-stimulated, followed by supernatant and cell collection after 24-hour incubation. Supernatant was then used for cytokine/chemokine ELISA and inserted into the bottom compartment of a TransWell system utilizing mTHP-1 cells in the upper compartment for cell migration assay, followed by evaluation with flow cytometry. Collected cells were used for cytokine/chemokine-coding mRNA detection using qRT-PCR. **b**. Proinflammatory cytokine expression in mTHP-1 and MTHP-1 cells transduced with Wuhan and Omicron ORF9b. Highlighted borders indicate adjusted *p*-values < 0.05. **c**. Proinflammatory cytokine/chemokine mRNA expression in LPS-treated mTHP-1-ORF9b cells. **d**. Proinflammatory cytokine/chemokine concentrations in the supernatant of LPS-treated mTHP-1-ORF9b cells. Error bars represent 95% CI values. **e**. mTHP-1 cell migration via TransWell inserts in the presence of untreated (NO LPS), 100 ng/ml LPS-treated (LPS 100) and 100 ng/ml LPS- and anti-CCL2 IgG-treated (LPS 100 + Anti-CCL2) mTHP-1-ORF9b cell growth media. Cell movement expressed as percentage of CFSE+ cells present at the bottom compartment of the TransWell system. Results were normalized according to untreated control mTHP-1 cells. **f**. CCL2-associated migration via TransWell inserts in the presence of mTHP-1-ORF9b growth media. Impact of CCL2 was calculated by subtracting migrated cell amount in the presence of 100 ng/ml LPS- and anti-CCL2 IgG-treated cell growth media from the total migrated cell number in the presence of 100 ng/ml LPS-treated cell growth media. Cell movement expressed as percentage of CFSE+ cells present at the bottom compartment of the TransWell system. Results were normalized according to untreated control mTHP-1 cells. \**p*⍰< ⍰0.05, \*\**p*⍰< ⍰0.01, \*\*\**p*⍰< ⍰0.001, \*\*\*\**p*⍰< ⍰0.0001.

Downregulation of proinflammatory cytokine/chemokine-coding genes observed in mTHP-1-ORF3a was either retained in Wuhan ORF3a-transduced MTHP-1 cells or returned to baseline in Omicron ORF3a-transduced MTHP-1 cells (Fig. 6A). LPS treatment of mTHP-1-ORF3a cells demonstrated a varied response largely similar to baseline mRNA expression levels of proinflammatory cytokines (Fig. 6B). Combined data from both LPS concentrations revealed a significant difference of *CCL2, CXCL11* and *IL1B* mRNA expression between Wuhan and Omicron ORF3a-transduced cells, demonstrating upregulation in Omicron ORF3a-transduced mTHP-1 cells (Fig. 6B). These findings are in line with alterations observed in cytokine/chemokine production, where Wuhan ORF3a-transduced mTHP-1 cells demonstrated decreased CCL2, CCL4 and IL-1β concentrations compared to control cells following LPS treatment (Fig. 6C). While Omicron ORF3a-transduced mTHP-1 cells exhibited decrease of CCL2 and IL-1β production, similar to Wuhan ORF3a-transduced mTHP-1 cells, CCL4 and CXCL11 concentrations were comparable to the baseline production following LPS treatment (Fig. 6C). As expected, both LPS-untreated control and mTHP-1-ORF3a cells exhibited marginal proinflammatory cytokine/chemokine mRNA expression and production (Supplementary Fig. 5B). In line with cytokine mRNA expression and production results, mTHP-1 cell movement through a Transwell membrane was significantly decreased in the presence of LPS-treated Wuhan mTHP-1-ORF3a growth media (Fig. 6D). A significant CCL2-associated movement decrease was observed in the presence of LPS-treated Wuhan mTHP-1-ORF3a growth media (Fig. 5E). In combination, this data suggests a persistent downregulation of proinflammatory cytokine/chemokine production in Wuhan and Omicron ORF3a-transduced mTHP-1 cells in the presence of TLR4 stimulation.

**Fig. 6.**
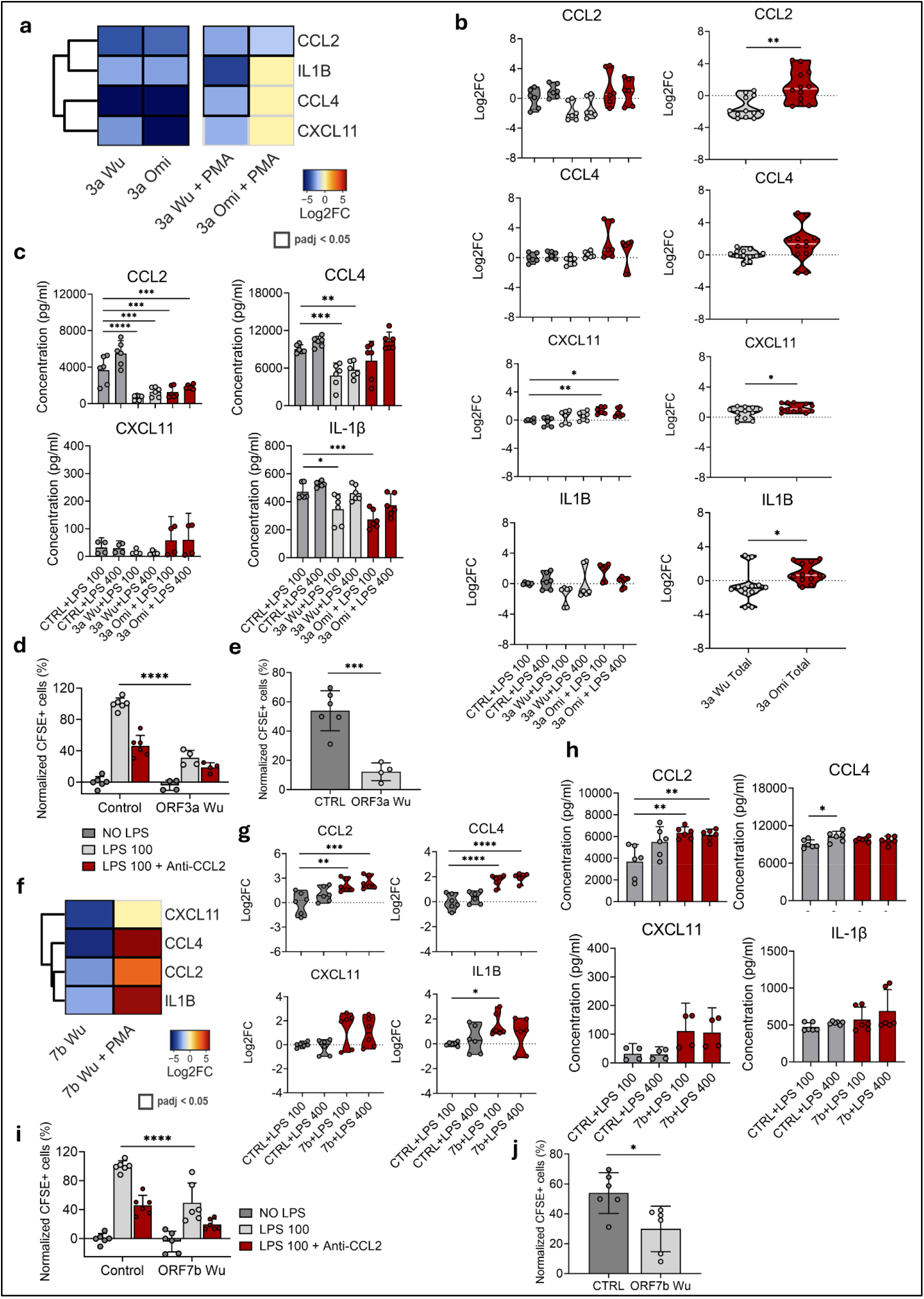
Proinflammatory cytokine/chemokine expression and production changes in ORF3a expressing mTHP-1 cells following TLR4 stimulation. **a**. Proinflammatory cytokine expression in mTHP-1 and MTHP-1 cells transduced with Wuhan and Omicron ORF3a. Highlighted borders indicate adjusted *p*-values < 0.05. **b**. Proinflammatory cytokine/chemokine mRNA expression in LPS-treated mTHP-1-ORF3a cells. **c**. Proinflammatory cytokine/chemokine concentrations in the supernatant of LPS-treated mTHP-1-ORF3a cells. Error bars represent 95% CI values **d**. mTHP-1 cell migration via TransWell inserts in the presence of untreated (NO LPS), 100 ng/ml LPS-treated (LPS 100) and 100 ng/ml LPS- and anti-CCL2 IgG-treated (LPS 100 + Anti-CCL2) mTHP-1-ORF3a cell growth media. Cell movement expressed as percentage of CFSE+ cells present at the bottom compartment of the TransWell system. Results were normalized according to untreated control mTHP-1 cells. **e**. CCL2-associated migration via TransWell inserts in the presence of mTHP-1-ORF3a growth media. Impact of CCL2 was calculated by subtracting migrated cell amount in the presence of 100 ng/ml LPS- and anti-CCL2 IgG-treated cell growth media from the total migrated cell number in the presence of 100 ng/ml LPS-treated cell growth media. Cell movement expressed as percentage of CFSE+ cells present at the bottom compartment of the TransWell system. Results were normalized according to untreated control mTHP-1 cells. **f**. Proinflammatory cytokine expression in mTHP-1 and MTHP-1 cells transduced with ORF7b. **g**. Proinflammatory cytokine/chemokine mRNA expression in LPS-treated mTHP-1-ORF7b cells. **h**. Proinflammatory cytokine/chemokine concentrations in the supernatant of LPS-treated mTHP-1-ORF7b cells. Error bars represent 95% CI values. **i**. mTHP-1 cell migration via TransWell inserts in the presence of mTHP-1-ORF7b cell growth media (see description of panel E). **j**. CCL2-associated migration via TransWell inserts in the presence of mTHP-1-ORF7b growth media (see description of panel F). \**p*⍰< ⍰0.05, \*\**p*⍰< ⍰0.01, \*\*\**p*⍰< ⍰0.001, \*\*\*\**p*⍰< ⍰0.0001.

Significant upregulation of proinflammatory cytokine/chemokine-coding gene expression was observed in MTHP-1-ORF7b cells, sharply contrasting the downregulation of *CXCL11, CCL4, CCL2* and *IL1B* in mTHP-1-ORF7b cells (Fig. 6F). LPS treatment resulted in upregulation of *CCL2, CCL4* and *IL1B* expression in 100 ng/ml LPS-treated mTHP-1-ORF7b cells (Fig. 6G). Cytokine/chemokine production in LPS-treated mTHP-1-ORF7b cells appeared largely unchanged compared to the LPS-treated control cells (except for CCL2 production which demonstrated a marked increase) (Fig. 6H). Interestingly, mTHP-1 cell movement through a Transwell membrane demonstrated a decrease in the presence of LPS-treated mTHP-1-ORF7b growth media (Fig. 6I). However, CCL2-associated movement decrease was marginally significant (Fig. 6J), largely consistent with results demonstrating return to baseline LPS-associated expression of proinflammatory cytokines/chemokines in mTHP-1-ORF7b cells.

### SARS-CoV-2 ORF3a and ORF9b may downregulate inflammatory response-related genes characteristic of surviving Omicron patients

In order to identify putative SARS-CoV-2 AP roles in COVID-19 patients, RNAseq datasets of BAL cells from patients infected with SARS-CoV-2 Omicron lineages and characterized by severe pneumonia were used for comparison (44). DEGs of COVID-19 survivor and non-survivor BAL samples were compared while Hospital Acquired Pneumonia/Community-Acquired Pneumonia (HAP/CAP) patient datasets served as controls. Gene set enrichment analysis (GSEA) revealed that all most significantly altered pathways in non-survivor vs control and survivor vs control gene sets were upregulated, while survivor vs non-survivor gene sets demonstrated downregulation, similar to observations in mTHP-1-ORFs (Fig. 8A). Most significantly altered Gene Ontology Biological Process (GOBP) pathways were related to immune response, such as *GOBP Phagocytosis, GOBP Myeloid Cell Activation Involved In Immune Response, GOBP Regulation Of Viral Genome Replication* and *GOBP Neutrophil Migration* (Fig. 8A). Similar to mTHP-1-ORFs (Fig. 2A), highly significant alterations were observed in inflammatory response-related pathways, such as *GOBP Inflammatory Response* and *GOBP Positive Regulation of Cytokine Production*, demonstrating significant number of downregulated terms in COVID-19 survivor group when compared to the non-survivor group (*GOBP Inflammatory Response* NES=-2.014, FDR≈0), and mTHP-1-ORF3a and - ORF9b compared to mTHP-1 WT cells of Wuhan (*GOBP Inflammatory Response* NES=-1.9, FDR≈0 and NES=-2.066, FDR≈0, respectively) and Omicron (*GOBP Inflammatory Response* NES=-1.879, FDR≈0 and NES=-2.426, FDR≈0, respectively) origin (Fig. 8B).

Observing alterations in individual DEGs within the *GOBP Inflammatory Response* pathway, 52 genes were shared between COVID-19 survivors vs non-survivors, and SARS-CoV-2 Omicron ORF3a- and ORF9b-transduced mTHP-1 cells (Fig. 8C). Majority of the most significantly downregulated genes in mTHP-1-ORF3a and -ORF9b cells, shared with COVID-19 survivors vs non-survivors, were the terminal inflammatory response genes, such as *CCL4, MMP9, C5AR1, CXCL8, CCL2, TNF, CXCL2, CCL3* and *IL1B*, expressed downstream of NF-κB pathway (Fig. 8D), with *MMP9, C5AR1, CXCL8, CXCL2* and *CCL3* demonstrating significant downregulation in COVID-19 survivor vs non-survivor dataset as well, with *CCL4* and *IL1B* approaching significance (adj *p* = 0.06, adj *p* = 0.07, respectively). Interestingly, mTHP-1 cells transduced with Omicron ORF3a exhibited stronger downregulation for all terminal inflammatory response genes, except for *TNF*, compared to their Wuhan ORF3a-transduced counterparts, while *CXCL2* was downregulated only in Omicron ORF3a- and ORF9b-transduced cells.

Comparatively less pronounced downregulation can be observed along the canonical IKKα/β/γ-NF-κB signaling pathway axis in both COVID-19 survivors vs non-survivors and mTHP-1-ORFs, with mild downregulation in NF-κB negative feedback loop signaling molecule IκBα-coding gene (*NFKBIA*)(Log2FC from -1.06 to -1.55 in mTHP-1-ORFs (adj *p* <0.05) and -1.84 in COVID-19 survivor vs control BAL (adj *p* = 0.12)) and phosphorylate NF-κB p65 amplifier *NFKBIZ* (Log2FC from -1.88 to -2.46 in mTHP-1-ORFs (adj *p* = 0.05) and Log2FC -1.75 in COVID-19 survivor vs control BAL (adj *p* = 0.08))(Fig. 8D). Moreover, NF-κB-related feed-forward amplifiers of inflammatory response, namely *JUN* and *CEBPB*, demonstrated mild downregulation in both COVID-19 survivors vs controls (Log2FC -1.22, adj *p* = 0.14 and Log2FC -2.07, adj *p* = 0.04, respectively) and mTHP-1-ORFs, albeit mostly in ORFs of Omicron origin. Additional downregulation has also been observed in TLR signaling-related genes, such as LY96, CD14 (TLR2/4 signaling) and TLR8 (Fig. 8D), with varying degree of significance between datasets.

Most notably, consistently significantly and robustly downregulated genes were identified along the type I IFN-signaling axis in COVID-19 survivors vs non-survivors and mTHP-1-ORFs, including *CXCL10, GBP2* and *CST7* (Fig. 8D). Strongest downregulation in all tested datasets was observed in *CXCL10* which demonstrated the most significant alteration in COVID-19 survivors vs non-survivors dataset (adj *p* = 0.003) and a notable increase in downregulation in Omicron ORF3a- and ORF9b-transduced mTHP-1 cells (Log2FC -10.14 and -6.15, respectively), compared to their Wuhan ORF-transduced counterparts (Log2FC -5.46 and -5.48, respectively). Both *GBP2* and *CST7* demonstrated significant downregulation in COVID-19 survivors vs non-survivors (Log2FC -2.08, adj *p* = 0.02 and Log2FC -3.07, adj *p* = 0.02, respectively) and mTHP-1-ORFs, with more robust downregulation levels observed in Omicron ORF3a- and ORF9b-transduced mTHP-1 cells.

## DISCUSSION

The continuous emergence of novel SARS-CoV-2 lineages, primarily belonging to the Omicron variant, together with the attenuation of symptoms in COVID-19 cases and growing concerns surrounding Long COVID have changed the landscape of SARS-CoV-2 infection compared with the early phase of the pandemic in 2019/2020. Although a substantial body of knowledge regarding SARS-CoV-2 biology has accumulated over the past years, the number of new mechanistic findings is gradually declining. The present study aims to address underexplored aspects of SARS-CoV-2 virology, specifically the interaction between APs and host immune cells, considering both the evolutionary importance of Omicron lineages and the role of myeloid cells in COVID-19 immunopathology. Recently, we identified metabolic and mitochondrial alterations associated with ORF3a, ORF9b, ORF9c and ORF10 expression in A549 lung epithelial cells (38). In the present study, we shifted our focus towards the primary cells responsible for generating inflammatory response following SARS-CoV-2 infection – monocytes and macrophages – and constructed SARS-CoV-2 AP-expressing mTHP-1 cells to evaluate intracellular alterations caused by different SARS-CoV-2 ORFs. Using a multilayered approach employing transcriptomics coupled with functional validation (qRT-PCR, ELISA and TransWell cell migration assays) we identified ORF3a, ORF9b and ORF7b APs that most prominently alter immune response-related pathways. In particular, ORF3a and ORF9b expression was associated with pronounced downregulation of pro-inflammatory mediator genes and suppression of cytokine/chemokine production in mTHP-1 cells, an effect often exacerbated in cells expressing Omicron-derived variants of these proteins. Finally, several inflammatory response-related genes downregulated in mTHP-1-ORF3a and mTHP-1-ORF9b cells overlapped with genes reduced in BAL samples from Omicron-infected survivors compared with non-survivors.

Our initial comparison between A549-ORF and mTHP-1-ORF cells revealed substantially different transcriptomic pictures (Fig. 1). Interestingly, while multiple genes related to inflammatory response were mostly upregulated (e.g. *TLR4, IL1B*) in A549-ORF3a, -ORF7b and -ORF9b cells, most inflammatory and immune response genes were downregulated in mTHP-1 cells expressing these same ORFs (Supplementary Fig. 2 and 3). This indicates cell type-dependent alterations associated with SARS-CoV-2 AP expression. This is unsurprising considering differences in alterations between not only epithelial and myeloid cells (45), but different types of myeloid cells as well (46) in response to SARS-CoV-2 infection. Therefore, establishing the roles of individual viral proteins requires analysis of multiple cell systems associated with different outcomes relating to viral pathobiology. While A549 and other epithelial cell models are commonly employed for SARS-CoV-2 protein research due to lung epithelial cells being the primary targets for SARS-CoV-2 infection, THP-1 or similar myeloid cell models are less frequently included, despite their importance in generating the hyperinflammatory response related to severe COVID-19. Selection of putative targets for COVID-19 treatment therefore must take cell-specific response into account.

While multiple intracellular targets of SARS-CoV-2 AP have previously been identified, ORF and immune response-related factor interactions are of particular interest, demonstrating the role of some APs as putative immunomodulators. ORF3a, ORF7b and ORF9b have all been characterized as exhibiting immunomodulatory capacity, associated with IFN signaling (13,18,20) and inflammatory response induction (9,34) or suppression (13,14,21). Alterations in canonical NF-κB signaling pathway are responsible for majority of inflammatory response modulations associated with ORF3a (direct interaction between ORF3a and both IKKβ and NEMO resulting in enhanced NF-κB activity)(9), ORF7b (direct interaction with MAVS and inhibition of RIG-I-like receptor signaling pathway)(21), and ORF9b (inhibition of IKKα signaling via NEMO interaction, resulting in reduced NF-κB nuclear translocation)(18) expression. In turn, cytokine/chemokine expression and production patterns are dysregulated in various ways. Our findings demonstrate changes in the transcriptomic landscape of ORF3a-, ORF7b- and ORF9b-expressing myeloid cells, with most significantly altered foci upstream and downstream of NF-κB, in addition to important IFN signaling pathway terms (Fig. 3B). While mTHP-1-ORF3a, -ORF7b and -ORF9b all demonstrated widespread immune response-related gene downregulation, further experiments in combination with previous findings revealed more nuanced mechanisms specific to particular ORFs.

While multiple studies have identified ORF3a as capable of priming NLRP3 inflammasome and promoting cytokine and chemokine release via NF-κB pathway (9), interestingly, our study demonstrates not only a significant downregulation of inflammatory response-related genes (e.g. *CCL4, CXCL11, CCL2, TNF, IL1B*) in ORF3a-transduced mTHP-1 and MTHP-1 cells (Fig. 3B, E), but dampened cytokine/chemokine (IL-1β, CCL2, CCL4) production (Fig. 6C) and suppressed monocyte response to mTHP-1-ORF3a growth media following LPS challenge as well (Fig. 6I). While NF-κB-mediated cytokine production increase in ORF3a expression systems have been identified in multiple studies (9,11,36,37,47), lower abundance of both p-NF-κB and NF-κB have been identified in ORF3a overexpressing cells, treated with poly(I:C)(13). Most recently, evidence suggesting complex dampening of both NF-κB pathway specifically and inflammatory response generally have appeared (48,49). Most previous studies investigating impact of ORF3a on host cells have focused on A549 and other epithelial cells, with very few studies investigating interactions between ORF3a and myeloid cells (9,34). Instead of a transiently transfected mTHP-1-ORF3a cells used in Ambrożek-Latecka et al., we have used stably expressing mTHP-1-ORF3a system, demonstrating significantly contrasting inflammatory response-related factor expression and production (34). While inflammasome activation and significant cytotoxicity in THP-1 cells may be observed soon after ORF3a expression, stabilized mTHP-1-ORF3a population may constitute surviving cells exhibiting immunoparalysis, characterized by downregulation of inflammatory response-related genes and blunted response to LPS stimulation. The concept of innate immune memory and similar tolerogenic recall responses have previously been described, most notably illustrated by dampened inflammatory responses in monocytes of sepsis-recovered patients (50). Innate immune memory associated with SARS-CoV-2 infection has previously been demonstrated in murine model, exhibiting pro-inflammatory cytokine/chemokine (including IL-1β and CCL4) reduction as a response to influenza A/PR/8/34 infection in SARS-CoV-2-recovered animals (51). However, no studies have attempted to investigate myeloid cell response to TLR stimulation in SARS-CoV-2 protein-expressing cells to date. While few studies to date have compared the differences between the impact of SARS-CoV-2 APs of different VOI origin on cellular processes, it has previously been suggested that ORF3a of Omicron variants harboring T223I mutation are characterized by decrease in lipid droplet accumulation and reduced ORF3a-Vps39 interaction in transduced Caco-2 cells (52). While similar studies in myeloid cells are yet to be repeated, lipid droplet stores in macrophages serve as precursors for pro-inflammatory mediator (e.g. prostaglandin-E2, IL-6) production (53) and contribute to monocyte to macrophage differentiation (54). Moreover, reduced ORF3a-Vps39 interaction may result in enhanced autophagy and in turn suppress NLRP3 inflammasome activation (55). Our findings are therefore in line with reduced acute inflammation seen in Omicron patients and findings demonstrating associations between Omicron-related ORF3a mutations and reduced NLRP3 inflammasome and pro-inflammatory mediator production (52,55). Consequently, the effect of ORF3a in myeloid cells may be complex and temporally sensitive, demonstrating context-dependent activation and suppression of inflammatory response. Therefore, further experiments comparing transiently and stably ORF3a-transduced myeloid cells of both Wuhan and Omicron origin should be carried out to further explore complex immunomodulatory role of ORF3a.

Conversely to SARS-CoV-2 ORF3a, our findings from mTHP-1-ORF9b cells agree with most studies describing ORF9b as an immunosuppressive protein acting as both type I IFN and NF-κB signaling pathway antagonist (18,56,57). Contrary to mTHP-1-ORF3a, we have discovered that downregulation of inflammatory response-related genes in Wuhan ORF9b expressing THP-1 is at least partly reversible in MTHP-1 cells (Fig. 4D). Conversely, Omicron ORF9b expressing mTHP-1 cells not only exhibited a higher proportion of inflammatory response-related genes but retained a larger number of significantly downregulated genes in MTHP-1 cells compared to their Wuhan counterparts (Fig. 4E). This trend remains at a post-transcriptional level as well, with LPS triggering a significantly lower cytokine/chemokine response (Fig. 5D) and results in suppressed monocyte movement in LPS-stimulated mTHP-1-ORF3a growth media (Fig. 5E) compared to both control and Wuhan ORF9b expressing mTHP-1 cells. Since most previous studies have employed epithelial cell models for ORF9b investigation, our findings are difficult to directly compare. However, similarly altered terms can be observed in myeloid cells. Most notably, ORF9b interrupts K63-linked polyubiquitination of NEMO, inhibiting IKKα signaling and NF-κB nuclear translocation, resulting in both type I IFN and cytokine suppression (14). Considering that NF-κB signaling pathway is primarily altered by ORF9b in a post-transcriptional fashion, our findings are in line with the current model, demonstrating that ORF9b-associated inhibition of NF-κB nuclear translocation results in significant downregulation of cytokine/chemokine-coding genes (Fig. 3B), but not terms upstream of NF-κB. While transcriptomic profiles of both Wuhan and Omicron ORF9b expressing mTHP-1 cells are similar, stark difference can be observed in cytokine/chemokine production which may be related to accumulated ORF9b volume rather than Omicron mutation-related changes in cell transcriptome. Indeed, ORF9b nucleotide substitution (28271A>T), which has previously been suggested to facilitate a weak Kozak initiation, resulting in increased leaky scanning translation of ORF9b from the N sgRNA, leads to increased ORF9b accumulation in host cells, likely exacerbating ORF9b-associated dampening of inflammatory responses (30,58). Therefore, at least in myeloid cells, while TLR stimulation or an active state may return cells to baseline cytokine/chemokine signaling patterns in the presence of ORF9b, increased accumulation of the protein in the case of Omicron may arrest NF-κB signaling pathway in a dose-dependent fashion, resulting in an overall blunted inflammatory response, which, similar to ORF3a, is in line with reduced acute inflammation seen in Omicron patients. Overall, ORF9b interaction with host cell inflammatory response appears to be more straightforward, compared to ORF3a. However, further experiments investigating direct interactions between ORF9b and myeloid cell factors should be carried out in order to establish the immunomodulatory mechanism of ORF9b.

Similar to Wuhan ORF9b-transduced mTHP-1 cells, SARS-CoV-2 ORF7b-transduced mTHP-1 cells, while demonstrating significant downregulation of immune response-related genes (Fig. 3B), not only did not exhibit a markedly lowered cytokine/chemokine response following LPS stimulation, compared to control mTHP-1 cells (Fig. 6H), but demonstrated widespread return to baseline expression and an enhanced expression of some immune response-related genes in MTHP-1 cells, e.g. exhibiting Log2FC > 4 in *IL1B, CCL4, CXCL8* and *TNF* (Fig. 3D). As is the case with ORF9b, to our knowledge, no studies exploring ORF7b impact on myeloid cells are currently available making comparisons with already existing literature, primarily utilizing epithelial cell models, complicated. Currently available data suggests a complex role of ORF7b with regards to both IFN and cytokine production pathways, with multiple conflicting findings. While few studies suggest a RIG-I-like receptor signaling pathway inhibition and, in turn, IFNβ promotor activity suppression (20,21), another study found upregulation of *IFNB* mRNA and increased IFNβ production in ORF7b expressing cells (23). Similarly, while one study demonstrated increase in TNFα and IL6 production and gene expression in ORF7b expressing cells (23), another study proposed MAVS-RLR signaling pathway suppression, resulting in downregulated TNFα gene expression (21). Together, these data suggest a context-dependent effect of ORF7b in myeloid cells, potentially exhibiting IFN and inflammatory response-related pathway downregulation via interference with MAVS-TRAF6 binding in monocytes, as seen in epithelial cell models. However, previously suggested promotion of STAT1 nuclear translocation by ORF7b (23), while lacking impact on inflammatory response pathways in otherwise unaltered cells, may trigger a substantial upregulation of proinflammatory cytokines following TLR signaling, since translocated STAT1 is recruited directly into TLR signaling pathway where it mediates proinflammatory cytokine production. Therefore, while in epithelial cells ORF7b may primarily serve as type I IFN antagonist, in myeloid cells it may contribute to hyperinflammatory response in concordance with additional signaling (driven by e.g. SARS-CoV-2 Spike-TLR4 interaction). Interestingly, expression of *TLR8* – gene coding for a PRR responsible for intracellular RNA detection – not only did not return to baseline expression in MTHP-1-ORF7b cells but was downregulated further (Fig. 3D). This finding is in line with immune evasion role of ORF7b, suggesting employment of specific strategy directed at anti-viral response-related pathways, but not antigen recognition system generally. Overall, ORF7b role in myeloid cells appears complex, partly overlapping with findings from epithelial cells, but exhibiting unique and paradoxical behavior as well. Further studies employing myeloid cell models and examining transcriptomic and production changes coupled with various types of secondary signals (such as poly(I:C)) may help elucidate the mechanism of ORF7b at various states.

While mutations observed in AP-coding genes of the Omicron variants of SARS-CoV-2 are rather limited in scope, they have been suggested as putative contributors to viral adaptations and reduced pathogenicity seen in Omicron patients (7). Therefore, investigations aimed at examining the effect of APs in Omicron-infected hosts, especially comparing survivor and non-survivor data, would not only help determine the evolution of AP role, but evaluate their contribution to reduced pathogenicity and COVID-19 morbidity as well. A previous study has demonstrated downregulatory trends in many inflammatory response-related genes in BAL samples of Omicron patients who survived the infection, compared to non-survivors (44). In order to compare transcriptomes of Omicron survivors with *in vitro* data from mTHP-1-ORF cells and identify similarly DE terms in both datasets, we have acquired transcriptomic data utilized in Wang et al. (44). SARS-CoV-2 patient transcriptomes demonstrate that non-survivor group of patients exhibit upregulation of pathways related to inflammatory pathway response, including upregulation of chemokines (e.g. *CCL2, CCL4*), cytokines (e.g. *IL1B*) and other pro-inflammatory factors (44)(Fig. 7A), suggesting a correlation between increased mortality from COVID-19 and hyperinflammation (59). Conversely, SARS-CoV-2 survivors exhibited downregulation of multiple inflammatory response-related pathways and genes, including *CCL2, CCL4* or *IL1B* (Fig. 7A). Incidentally, downregulation of inflammatory response pathway and genes such as *CCL2, CCL4, CCL3, IL1B, TNF* and *CXCL11* was observed in mTHP-1-ORF3a, -ORF9b and -ORF7b cells (Fig. 3B), demonstrating putative inflammatory response antagonism of ORF3a, ORF9b and ORF7b on a transcriptomic level. Furthermore, increase in downregulation associated with ORF3a and ORF9b of Omicron origin suggests a link between these APs and attenuated inflammatory response in Omicron patients.

**Fig. 7.**
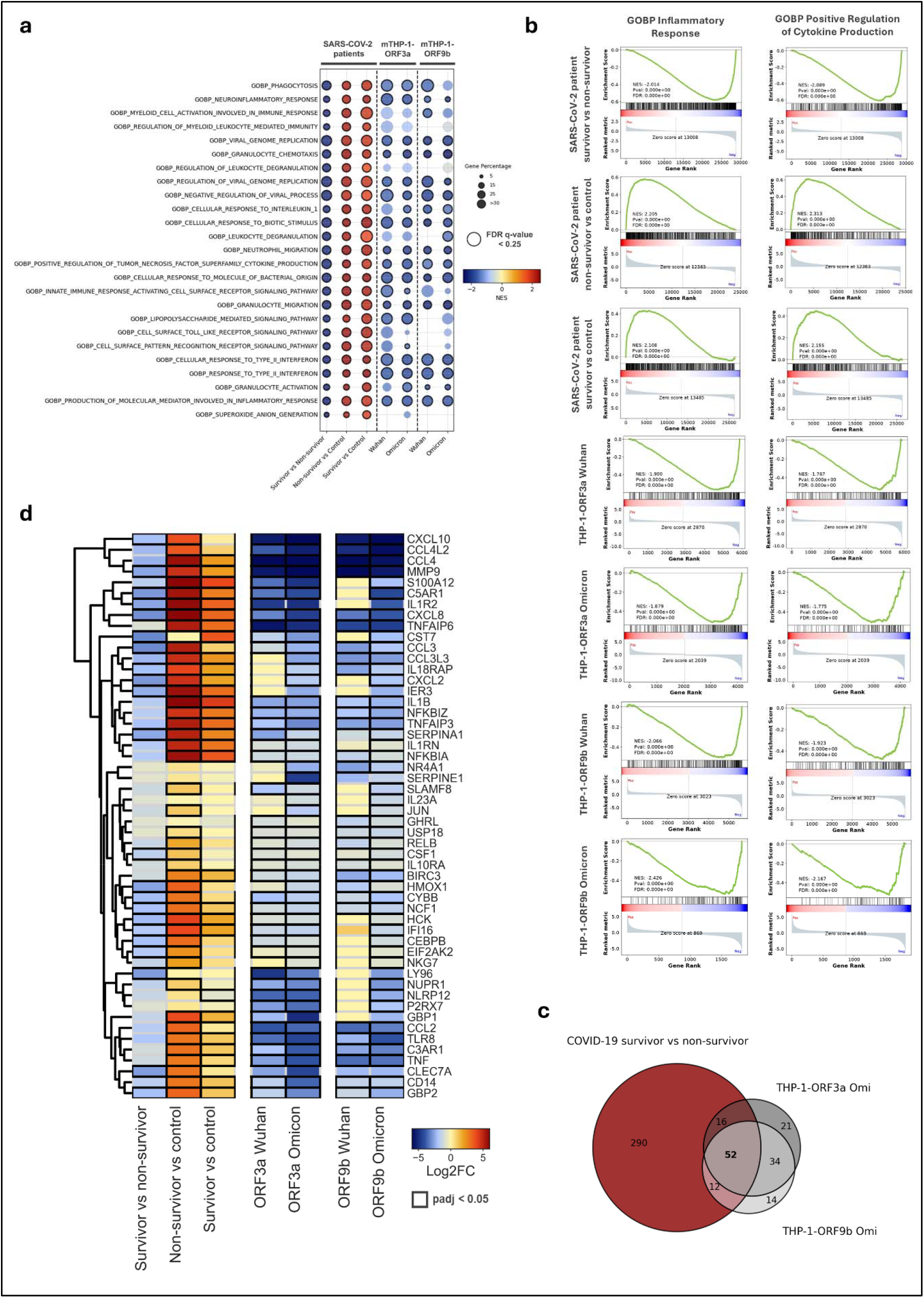
Comparison between transcriptomic alterations of SARS-CoV-2 Omicron patient BAL samples cells and ORF9b- and ORF3a-transduced mTHP-1 cells. **a**. Dot plot representing SARS-CoV-2 Omicron patient BAL, mTHP-1-ORF3a and -ORF9b normalized enrichment score (NES) distribution according to the 25 most significantly altered Gene Ontology Biological Process (GOBP) pathways based on gene expression in BAL between survivors and non-survivors of Omicron infection. The size and color of each dot represents the altered gene percentage in a given pathway and NES level, respectively. Dots with highlighted borders represent data points with multiple false discovery rate (FDR) q-values of <0.25. **b**. Gene Set Enrichment Analysis plots of the *GOBP Inflammatory Response* and *GOBP Positive Regulation of Cytokine Production* hallmark gene sets for SARS-CoV-2 Omicron patient BAL and mTHP-1-ORF3a and - ORF9b samples. **c**. Venn diagram demonstrating overlap of DEGs belonging to *GOBP Inflammatory Response* hallmark gene set between COVID-19 survivor vs non-survivor, mTHP-1-ORF3a and -ORF9b datasets. **d**. Heatmap depicting differential expression profile (based on Log2-fold change) of 52 overlapping genes from C in SARS-CoV-2 Omicron patient BAL, mTHP-1-ORF3a and -ORF9b samples. Highlighted borders represent data with adjusted *p*-values < 0.05.

While our study establishes immune response alterations associated with SARS-CoV-2 ORF3a, ORF7b and ORF9b expression in myeloid cells, the following limitations need to be addressed. Firstly, although cell line models, such as A549 and THP-1, are routinely used to research SARS-CoV-2 pathobiology, findings acquired from immortalized cell lines need to be validated in primary cells in order to be attributable to physiological conditions *in vivo*. Moreover, while our study employed mTHP-1 and MTHP-1 cells, myeloid cells of different subtypes not only produce a markedly different response to SARS-CoV-2 infection (60), but respond differently depending on the co-culture environment as well. Secondly, direct comparison between Omicron patient BAL and *in vitro* mTHP-1-ORF transcriptomics data is difficult due to the following factors. Firstly, while cells retrieved from BAL samples likely contain a population of alveolar macrophages, their mRNAs are not analyzed separately from other cell mRNAs, resulting in metatranscriptome vs transcriptome dataset comparisons. Secondly, comparatively few Omicron patient samples met our data quality criteria, resulting in a limited viable sample size. In turn, significantly lower amount of genes demonstrated adjusted *p*-values < 0.05, compared to the original study (44). Finally, Omicron patient transcriptomics data represents alterations in BAL sample cells resulting from full infection, rather than individual viral protein transduction. In combination, these factors limit the translational power of Omicron patient and *in vitro* data comparison. However, it provides a framework for future studies aiming for a more accurate analysis, such as transiently ORF-transduced human primary myeloid cell comparisons with Omicron patient myeloid cell data. Finally, while transcriptomics data provides valuable insights into mRNA expression landscape within SARS-CoV-2 ORF-transduced cells, post-transcriptional alteration evaluation requires protein-protein interactome mapping, employing proteomic or phosphoproteomic approaches. According to previous studies focusing on epithelial cells, most significant alterations carried out by SARS-CoV-2 APs are post-transcriptional in nature, rarely demonstrating DE of their target-coding genes, as suggested by our findings from myeloid cells as well. Therefore, while transcriptomic approach helps in providing an overview of the effect a specific ORF may have on cellular functioning at large, establishment of a more complete picture of SARS-CoV-2 ORF mechanism of action within host myeloid cells and differences exhibited by proteins of various VOIs requires further experiments investigating direct interactions between viral ORFs and host factors.

In conclusion, our findings include the first comprehensive investigation of transcriptomic changes observed in SARS-CoV-2 AP-transduced monocyte/macrophage cells, with ORF3a, ORF9b and ORF7b expression resulting in the most significant alterations associated with inflammatory response pathways. We have identified a significant downregulation of proinflammatory cytokine/chemokine-coding genes in mTHP-1 cells across all three ORFs, but various levels of impact associated with transcription of these same genes in MTHP-1 cells. Further experiments allowed us to reveal nuanced mechanisms by which these ORFs impact cytokine/chemokine expression and production, most notably associated with attenuated inflammatory response factors in Omicron ORF3a and ORF9b expressing mTHP-1 cells in the presence of TLR4 signaling, corresponding to reduced pathogenicity and inflammatory responses seen in Omicron-infected patients. Finally, we have identified inflammatory response-related genes downregulated in both Omicron survivors and ORF9b/ORF3a expressing mTHP-1 cells, suggesting possible contribution of these ORFs in Omicron cases characterized by reduced morbidity and mortality.

## MATERIALS AND METHODS

### 1. Cell culture and lentiviral transduction

A549 pulmonary epithelial cells (ATCC CRM-CCL-185) and THP-1 monocytic leukemia cells (ATCC TIB-202) were cultured in Dulbecco’s Modified Eagle Medium (DMEM) and Roswell Park Memorial Institute medium (RPMI), respectively, supplemented with 10% (v/v) heat-inactivated fetal bovine serum (FBS) 1% Penicillin-Streptomycin (100U/ml) and Amphotericin B. Cells were passaged at 80% confluence and 8 ×⍰10^5^ cells/ml concentration, periodically monitoring for mycoplasma contamination. All cells were cultured at 37°C in a 5% CO_2_, 90% humidity atmosphere.

Lentivirus particles containing the coding sequences of ORF3a, ORF6, ORF7a, ORF7b, ORF8, ORF9b, ORF9c, ORF8 and ORF10 of SARS-CoV-2 Wuhan-Hu-1 isolate and ORF3a and ORF9b of SARS-CoV-2 Omicron 21L (BA.2) isolate were generate at the Centro Nacional de Investigaciones Cardiovasculares (CNIC) Viral Vector Unit, following the previously described protocol (61). All sequences were codon-optimized for mammalian expression and were C-terminally 2 × Strep-tagged in their N-terminal or C-terminal domain to allow protein identification. For transduction of A549 cells, 1 ×⍰10^5^ cells were incubated with lentiviral vector at a MOI of 10 with 5 μg/ml polybrene for 24h in DMEM without puromycin, followed by 2 μg/ml puromycin treatment for successfully transduced cell enrichment. For transduction of THP-1 cells, double spinoculation protocol was used. Firstly, 1 ×⍰10^6^ cells were incubated with lentiviral vector at a MOI of 150 in the presence of 8 μg/ml polybrene for 48h in RPMI without puromycin. The process was repeated utilizing 24h incubation. Further transduced cell maintenance was carried out as outlined above with 2 μg/ml puromycin added to the culture media for successfully transduced cell selection. Both A549 and THP-1 control cells transduced with mock lentiviral vector and tagged with 2xStrep tag were also generated.

### 2. Differentiation of mTHP-1 cells into MTHP-1 cells

For differentiation into MTHP-1 cells, 1 ×⍰10^5^ cells/ml mTHP-1 cells were collected, centrifuged and resuspended in RPMI media containing PMA at concentration of 25 ng/ml. Cells were then transferred to cell culture flasks at 5 ×⍰10^5^ cells/flask and incubated for 48 hours at 37°C in a 5% CO_2_, 90% humidity. Finally, attachment of the cells to the bottom of the flask was observed, signifying successful differentiation, followed by replacement of maintenance media with RPMI without PMA and an additional 24-hour incubation.

### 3. RNA isolation and RNAseq of A549-ORF, mTHP-1-ORF and MTHP-1-ORF cells

Vector- and SARS-CoV-2 ORF-transduced A549, mTHP-1 and MTHP-1 cells were selected for RNA purification and sequencing as previously described (62). Sequence raw data quality control and filtering was performed as previously described (62). Genes for which the sum of raw counts across all of the samples was <10 were discarded. Differential expression and analysis was performed using DESeq2 1.46.0 R package (63).

### 4. LPS stimulation of mTHP-1 cells

For LPS stimulation experiments, control cells and mTHP-1 cells, transduced with ORF3a and ORF9b of Wuhan and Omicron origin, ORF6 and ORF7b, were transferred to individual wells at concentrations of 5 ×⍰10^5^ cells/ml. Cells were then treated with LPS at final concentrations of 100 and 400 ng/ml or mock-stimulated with 1x Phosphate Buffered Saline (PBS) and incubated for 24 hours at 37°C in a 5% CO_2_, 90% humidity atmosphere. Cells were then transferred to 1.5 ml tubes, centrifuged at 130 x g for 5 min at room temperature. Supernatant was collected and stored at -20°C for ELISA or cell migration experiments. Cells were resuspended in 350 µl RTL buffer (Qiagen) and stored in -80°C for RNA purification.

### 5. Real Time qPCR

RNA of LPS- and mock-stimulated mTHP-1 cells was isolated as previously described (38). Reverse transcription of 500 ng total RNA samples was performed using NZY First-Strand cDNA Synthesis Kit (nzytech) following manufacturer’s instructions. In order to measure the relative gene expression of *CCL2, CCL4, CXCL11* and *IL-1β*, real-time PCR was performed using NZYSpeedy qPCR Green Master Mix (nzytech). *GDAPH* was used as a housekeeping gene. Primer pair sequences used are described in Supplementary Table 1. LightCycler 96 (Roche) was used for real-time PCR under the following conditions: 5 min at 95°C followed by 45 cycles of 20 s at 94°C, 30 s at 57℃. Melting curve was constructed to ensure specificity of each PCR product. Transcript levels, obtained using 2-ΔΔCt calculation method, were normalized to mTHP-1 cells untreated with LPS.

### 6. ELISA

Supernatants of LPS- or mock-stimulated control or mTHP-1-ORF cells were used for determination of proinflammatory cytokine/chemokine concentrations. Human CCL2 (MCP-1), CCL4 (MIP-1b) and CXCL11 (I-TAC) Quantikine ELISA kits (R&D Systems) and IL-1β DuoSet ELISA kit (R&D Systems) were used according to manufacturer’s instructions. Optical density for each well was determined using Multiskan™ Sky Microplate Spectrophotometer (ThermoFisher Scientific) set to 450 nm and wavelength correction set to 540 nm. Concentrations of tested substances were calculated according to the standard curve.

### 7. Cell Migration Assay

TransWell plates with 5 µm pore-size inserts (Sigma-Aldrich) were used for evaluation of mTHP-1 cell migration. 600 µl of supernatant from unstimulated and LPS-stimulated mTHP-1-ORFs were added to the lower compartment of the system. Unstimulated control mTHP-1 cells at 1 ×⍰10^6^ cells/ml concentration were first labelled with 5 μM carboxyfluorescein succinimidyl ester (CFSE) and incubated for 20 min at 37 ºC. Subsequently, 3 ×⍰10^5^ cells/ml of labelled control mTHP-1 cells were transferred to the upper compartment of the TransWell system and incubated for 24 h at 37°C in a 5% CO_2_, 90% humidity atmosphere.

Migrated cells in the lower compartment were then quantified using a CytoFLEX flow cytometer (FITC channel; Beckman Coulter). To assess CCL2-dependent migration, 1 μg/mL anti-CCL2 (R&D Systems) was added to an additional LPS-stimulated well, while an isotype-matched antibody (IgG2B)(R&D Systems) served as negative control. Antibodies were pre-incubated with cells for 30 min at 37ºC with 5% CO_2_, and migration of labelled cells were also evaluated in these antibody-treated supernatants. CCL2-specific migration was calculated by subtracting the number of migrated cells in anti-CCL2-treated wells from those in LPS-only conditions. Finally, CCL2-dependent migration values in the supernatants of cells overexpressing selected SARS-CoV-2 APs were normalized to those of control cells.

### 8. COVID-19 patient BAL sample data acquisition and transcriptomics

Reads were obtained from Wang et al. (44), comprising 63 COVID-19 patients and 27 invasive ventilated HAP/CAP patients, and downloaded from the National Microbiology Data Center (Beijing Institute of Microbiology, Chinese Academy of Sciences). All raw reads were processed through the nf-core/rnaseq pipeline (release 3.18.0) (64) built with Nextflow 24.10.4 (65) with default settings, which encompasses adapter trimming, quality filtering, alignment, quantification, and generation of summary metrics. Specifically, adapter removal and quality trimming were performed with Trim Galore 0.6.10, aligned with STAR 2.7.11b (66) and quantified transcripts abundance with Salmon 1.10.3 (67) in alignment-based mode. Samples were aligned against reference genome GRCh38, and the gtf file from the same release was used for the transcripts. Bias-corrected counts without an offset were obtained from the pipeline usingrximport1.20.1 (68) with length scaling and imported into R 4.4.2 for downstream analysis. A subset of 12 highest-quality samples per condition were selected based on the lowest duplication rates according to Picard-tools Mark Duplicates 3.1.1 and the highest alignment percentages. Genes with <10 counts across all samples were discarded. DESeq2 1.46.0 (63) R package was used to obtain DEGs, with the design formula ∼condition, employing Benjamini–Hochberg correction and a significance cutoff of adjusted *p* < 0.05.

### 9. Statistical Analysis

RNAseq data analysis was performed according to procedures outlined in the transcriptomic sections above. For hierarchical clustering, VST-transformed expression values were averaged across replicates within each condition. Euclidean distances were calculated between condition-level expression profiles, and hierarchical clustering was performed using complete linkage (hclust). Metascape Gene List Analysis tool with Multiple Gene List default setting was used for Pathway Enrichment Analysis of all mTHP-1-ORF cells (39), while Protein-protein Interaction Enrichment Analysis was carried out using Molecular Complex Detection (MCODE) algorithm in Metascape. Additional Gene Set Enrichment Analysis (GSEA) of human patient BAL, mTHP-1-ORF3a and mTHP-1-ORF9b cell data was performed using GSEApy 1.1.11 library in Python, utilizing 1000 random permutations for FDR q value and normalized enrichment score (NES) acquisition.

Relative RNA expression, cytokine/chemokine concentration and CFSE+ cell number comparisons were performed using One-way ANOVA with Dunnett’s multiple comparisons test. Combined RNA comparisons between groups were carried out using unpaired t-test. Value of *p* < 0.05 were considered statistically significant.

### 10. Result Visualization

Heatmaps, ridgeline plots, venn diagrams and dot plots were constructed in Python 3.15 utilizing pandas, seaborn, matplotlib, numpy, matplotlib_venn, joypy and mygene libraries, unless otherwise stated in the figure description. Violin and bar plots were constructed using GraphPad Prism 10.3.0 (Dotmatics). Linear dendrograms were generated and customized using the dendextend package (69).

## DATA AVAILABILITY

All data are available in the main text or the supplementary materials. Source data are provided with this paper.

## ACKNOWLEDGEMENTS

This research work was funded several funding agencies, including Research Council of Lithuania (LMTLT), agreement No. S-PD-24-110, European Commission NextGenerationEU (Regulation EU 2020/2094), through CSIC’s Global Health Platform (PTI+ Salud Global) (COVID-19-117 and SGL2103015), Junta de Andalucía (CV20-20089), Spanish Ministry of Science projects (PID2021-123399OB-I00, CNS2023-145079 and PID2024-162356OB-I00). T.G.G is recipient of a Ramón y Cajal contract (RYC2021-031614-I) funded by MCIN/AEU/10.13039/ 501100011033 and NextGeneration EU/PRTR.

## AUTHOR CONTRIBUTIONS

M.M. led the study. M.M., I.K.K. and J.J.G. supervised the study. J.G., M.P-B., M.M., I.K.K., T.G.G. and B.D.L-A. designed the experiments. J.G., M.P-B., B.D.L-A., A.L.R., T.G.G., R.F.R. and L.M.G. performed the experiments. J.G., M.P-D., U.M.H., A.P. and A.S. analyzed the data. J.G., U.M.H., M.P-B. and A.P. performed data visualizations. J.G., M.M., I.K.K., J.J.G. and A.S. provided resources and reagents. M.M., I.K.K. and J.G. acquired funding. J.G., M.P-B., M.M. and I.K.K. wrote the manuscript. All authors reviewed and approved the manuscript.

## REFERENCES

1. Vora SM, Lieberman J, Wu H. Inflammasome activation at the crux of severe COVID-19. Nat Rev Immunol. 2021 Nov;21(11):694–703. doi:10.1038/s41577-021-00588-x

2. Mohandas S, Jagannathan P, Henrich TJ, Sherif ZA, Bime C, Quinlan E, et al. Immune mechanisms underlying COVID-19 pathology and post-acute sequelae of SARS-CoV-2 infection (PASC). Iqbal J, Zaidi M, editors. eLife. 2023 May 26;12:e86014. doi:10.7554/eLife.86014

3. Hoenigsperger H, Sivarajan R, Sparrer KM. Differences and similarities between innate immune evasion strategies of human coronaviruses. Curr Opin Microbiol. 2024 Jun 1;79:102466. doi:10.1016/j.mib.2024.102466

4. Tsukalov I, Sánchez-Cerrillo I, Rajas O, Avalos E, Iturricastillo G, Esparcia L, et al. NFκB and NLRP3/NLRC4 inflammasomes regulate differentiation, activation and functional properties of monocytes in response to distinct SARS-CoV-2 proteins. Nat Commun. 2024 Mar 7;15(1):2100. doi:10.1038/s41467-024-46322-8

5. Shahbaz S, Bozorgmehr N, Lu J, Osman M, Sligl W, Tyrrell DL, et al. Analysis of SARS-CoV-2 isolates, namely the Wuhan strain, Delta variant, and Omicron variant, identifies differential immune profiles. Microbiol Spectr. 2023 Sep 7;11(5):e01256–23. doi:10.1128/spectrum.01256-23

6. Relan P, Motaze NV, Kothari K, Askie L, Waroux OLP de, Kerkhove MDV, et al. Severity and outcomes of Omicron variant of SARS-CoV-2 compared to Delta variant and severity of Omicron sublineages: a systematic review and metanalysis. BMJ Glob Health. 2023 Jul 6;8(7). doi:10.1136/bmjgh-2023-012328

7. Padilla-Blanco M, García-García T, Grigas J, López-Ayllón BD, Garrido JJ, Oliva MA, et al. Hidden players of COVID-19: the evolving roles of SARS-CoV-2 accessory proteins. Front Immunol. 2025 Nov 28;16. doi:10.3389/fimmu.2025.1726698

8. Ambrożek-Latecka M, Kozlowski P, Hoser G, Bandyszewska M, Hanusek K, Nowis D, et al. SARS-CoV-2 and its ORF3a, E and M viroporins activate inflammasome in human macrophages and induce of IL1α in pulmonary epithelial and endothelial cells. Cell Death Discov. 2024 Apr 25;10(1):191. doi:10.1038/s41420-024-01966-9

9. Nie Y, Mou L, Long Q, Deng D, Hu R, Cheng J, et al. SARS-CoV-2 ORF3a positively regulates NF-κB activity by enhancing IKKβ-NEMO interaction. Virus Res. 2023 Apr 15;328:199086. doi:10.1016/j.virusres.2023.199086

10. Zhang X, Yang Z, Pan T, Long X, Sun Q, Wang PH, et al. SARS-CoV-2 ORF3a induces RETREG1/FAM134B-dependent reticulophagy and triggers sequential ER stress and inflammatory responses during SARS-CoV-2 infection. Autophagy. 2022 Nov 2;18(11):2576–92. doi:10.1080/15548627.2022.2039992

11. Zhang J, Cruz-Cosme R, Zhang C, Liu D, Tang Q, Zhao RY. Endoplasmic reticulum-associated SARS-CoV-2 ORF3a elicits heightened cytopathic effects despite robust ER-associated degradation. mBio. 2023 Dec 11;15(1):e03030–23. doi:10.1128/mbio.03030-23

12. Su J, Shen S, Hu Y, Chen S, Cheng L, Cai Y, et al. SARS-CoV-2 ORF3a inhibits cGAS-STING-mediated autophagy flux and antiviral function. J Med Virol. 2023;95(1):e28175. doi:10.1002/jmv.28175

13. Lundrigan E, Toudic C, Pennock E, Pezacki JP. SARS-CoV-2 Protein Nsp9 Is Involved in Viral Evasion through Interactions with Innate Immune Pathways. ACS Omega. 2024 Jun 18;9(24):26428–38. doi:10.1021/acsomega.4c02631

14. Wu J, Shi Y, Pan X, Wu S, Hou R, Zhang Y, et al. SARS-CoV-2 ORF9b inhibits RIG-I-MAVS antiviral signaling by interrupting K63-linked ubiquitination of NEMO. Cell Rep. 2021 Feb 16;34(7). doi:10.1016/j.celrep.2021.108761

15. Han L, Zhuang M, Deng J, Zheng Y, Zhang J, Nan M, et al. SARS-CoV-2 ORF9b antagonizes type I and III interferons by targeting multiple components of the RIG-I/MDA-5–MAVS, TLR3–TRIF, and cGAS– STING signaling pathways. J Med Virol. 2021 Sep;93(9):5376–89. doi:10.1002/jmv.27050

16. Kouwaki T, Nishimura T, Wang G, Oshiumi H. RIG-I-Like Receptor-Mediated Recognition of Viral Genomic RNA of Severe Acute Respiratory Syndrome Coronavirus-2 and Viral Escape From the Host Innate Immune Responses. Front Immunol. 2021 Jun 25;12. doi:10.3389/fimmu.2021.700926

17. Zodda E, Pons M, DeMoya-Valenzuela N, Calvo-González C, Benítez-Rodríguez C, López-Ayllón BD, et al. Induction of the Inflammasome by the SARS-CoV-2 Accessory Protein ORF9b, Abrogated by Small-Molecule ORF9b Homodimerization Inhibitors. J Med Virol. 2025;97(2):e70145. doi:10.1002/jmv.70145

18. Han L, Zhuang M, Deng J, Zheng Y, Zhang J, Nan M, et al. SARS-CoV-2 ORF9b antagonizes type I and III interferons by targeting multiple components of the RIG-I/MDA-5–MAVS, TLR3–TRIF, and cGAS– STING signaling pathways. J Med Virol. 2021 Sep;93(9):5376–89. doi:10.1002/jmv.27050

19. Janevska M, Naessens E, Verhasselt B. Impact of SARS-CoV-2 Wuhan and Omicron Variant Proteins on Type I Interferon Response. Viruses. 2025 Apr;17(4):569. doi:10.3390/v17040569

20. Shemesh M, Aktepe TE, Deerain JM, McAuley JL, Audsley MD, David CT, et al. SARS-CoV-2 suppresses IFNβ production mediated by NSP1, 5, 6, 15, ORF6 and ORF7b but does not suppress the effects of added interferon. PLOS Pathog. 2021 Aug 26;17(8):e1009800. doi:10.1371/journal.ppat.1009800

21. Xiao X, Fu Y, You W, Huang C, Zeng F, Gu X, et al. Inhibition of the RLR signaling pathway by SARS-CoV-2 ORF7b is mediated by MAVS and abrogated by ORF7b-homologous interfering peptide. J Virol. 2024 Apr 4;98(5):e01573–23. doi:10.1128/jvi.01573-23

22. Stukalov A, Girault V, Grass V, Karayel O, Bergant V, Urban C, et al. Multilevel proteomics reveals host perturbations by SARS-CoV-2 and SARS-CoV. Nature. 2021 Jun;594(7862):246–52. doi:10.1038/s41586-021-03493-4

23. Yang R, Zhao Q, Rao J, Zeng F, Yuan S, Ji M, et al. SARS-CoV-2 Accessory Protein ORF7b Mediates Tumor Necrosis Factor-α-Induced Apoptosis in Cells. Front Microbiol. 2021 Aug 13;12. doi:10.3389/fmicb.2021.654709

24. Liu L, Zhang L, Hao X, Wang Y, Zhang X, Ge L, et al. Coronavirus envelope protein activates TMED10-mediated unconventional secretion of inflammatory factors. Nat Commun. 2024 Oct 8;15(1):8708. doi:10.1038/s41467-024-52818-0

25. Barh D, Tiwari S, Rodrigues Gomes LG, Ramalho Pinto CH, Andrade BS, Ahmad S, et al. SARS-CoV-2 Variants Show a Gradual Declining Pathogenicity and Pro-Inflammatory Cytokine Stimulation, an Increasing Antigenic and Anti-Inflammatory Cytokine Induction, and Rising Structural Protein Instability: A Minimal Number Genome-Based Approach. Inflammation. 2023 Feb 1;46(1):297–312. doi:10.1007/s10753-022-01734-w

26. Wang W, Qu Y, Wang X, Xiao MZX, Fu J, Chen L, et al. Genetic variety of ORF3a shapes SARS-CoV-2 fitness through modulation of lipid droplet. J Med Virol. 2023;95(3):e28630. doi:10.1002/jmv.28630

27. Hossain A, Akter S, Rashid AA, Khair S, Alam ASMRU. Unique mutations in SARS-CoV-2 Omicron subvariants’ non-spike proteins: Potential impacts on viral pathogenesis and host immune evasion. Microb Pathog. 2022 Sep;170:105699. doi:10.1016/j.micpath.2022.105699

28. Islam MdA, Shahi S, Marzan AA, Amin MR, Hasan MN, Hoque MN, et al. Variant-specific deleterious mutations in the SARS-CoV-2 genome reveal immune responses and potentials for prophylactic vaccine development. Front Pharmacol. 2023 Feb 7;14:1090717. doi:10.3389/fphar.2023.1090717

29. Zhang C, Gerzanich V, Cruz-Cosme R, Zhang J, Tsymbalyuk O, Tosun C, et al. SARS-CoV-2 ORF3a induces COVID-19-associated kidney injury through HMGB1-mediated cytokine production. mBio. 2024 Sep 30;15(11):e02308–24. doi:10.1128/mbio.02308-24

30. Caobi A, Saeed M. Upping the ante: enhanced expression of interferon-antagonizing ORF6 and ORF9b proteins by SARS-CoV-2 variants of concern. Curr Opin Microbiol. 2024 Jun 1;79:102454. doi:10.1016/j.mib.2024.102454

31. Janevska M, Naessens E, Verhasselt B. Impact of SARS-CoV-2 Wuhan and Omicron Variant Proteins on Type I Interferon Response. Viruses. 2025 Apr;17(4):569. doi:10.3390/v17040569

32. Mohandas S, Jagannathan P, Henrich TJ, Sherif ZA, Bime C, Quinlan E, et al. Immune mechanisms underlying COVID-19 pathology and post-acute sequelae of SARS-CoV-2 infection (PASC). Iqbal J, Zaidi M, editors. eLife. 2023 May 26;12:e86014. doi:10.7554/eLife.86014

33. Sahanic S, Hilbe R, Dünser C, Tymoszuk P, Löffler-Ragg J, Rieder D, et al. SARS-CoV-2 activates the TLR4/MyD88 pathway in human macrophages: A possible correlation with strong pro-inflammatory responses in severe COVID-19. Heliyon. 2023 Nov 1;9(11). doi:10.1016/j.heliyon.2023.e21893

34. Ambrozek-Latecka M, Kozlowski P, Hoser G, Bandyszewska M, Hanusek K, Nowis D, et al. SARS-CoV-2 and its ORF3a, E and M viroporins activate inflammasome in human macrophages and induce of IL-1α in pulmonary epithelial and endothelial cells. Cell Death Discov. 2024 Apr 25;10(1):191. doi:10.1038/s41420-024-01966-9

35. Paunovic V, Vucicevic L, Misirkic Marjanovic M, Perovic V, Ristic B, Bosnjak M, et al. Autophagy Receptor p62 Regulates SARS-CoV-2-Induced Inflammation in COVID-19. Cells. 2023 Jan;12(9):1282. doi:10.3390/cells12091282

36. Xu H, Akinyemi IA, Chitre SA, Loeb JC, Lednicky JA, McIntosh MT, et al. SARS-CoV-2 viroporin encoded by ORF3a triggers the NLRP3 inflammatory pathway. Virology. 2022 Mar 1;568:13–22. doi:10.1016/j.virol.2022.01.003

37. Ahmad F, Keshri V, Singh SK. ORF3a of SARS-CoV-2 modulates PI3K/AKT signaling in human lung epithelial cells via hsa-miR-155-5p. Int J Biol Macromol. 2024 May 1;268:131734. doi:10.1016/j.ijbiomac.2024.131734

38. López-Ayllón BD, Marin S, Fernández MF, García-García T, Fernández-Rodríguez R, de Lucas-Rius A, et al. Metabolic and mitochondria alterations induced by SARS-CoV-2 accessory proteins ORF3a, ORF9b, ORF9c and ORF10. J Med Virol. 2024;96(7):e29752. doi:10.1002/jmv.29752

39. Zhou Y, Zhou B, Pache L, Chang M, Khodabakhshi AH, Tanaseichuk O, et al. Metascape provides a biologist-oriented resource for the analysis of systems-level datasets. Nat Commun. 2019 Apr 3;10(1):1523. doi:10.1038/s41467-019-09234-6

40. Junqueira C, Crespo Â, Ranjbar S, de Lacerda LB, Lewandrowski M, Ingber J, et al. FcγR-mediated SARS-CoV-2 infection of monocytes activates inflammation. Nature. 2022 Jun;606(7914):576–84. doi:10.1038/s41586-022-04702-4

41. Suzuki R, Yamasoba D, Kimura I, Wang L, Kishimoto M, Ito J, et al. Attenuated fusogenicity and pathogenicity of SARS-CoV-2 Omicron variant. Nature. 2022 Mar;603(7902):700–5. doi:10.1038/s41586-022-04462-1

42. Wang W, Qu Y, Wang X, Xiao MZX, Fu J, Chen L, et al. Genetic variety of ORF3a shapes SARS-CoV-2 fitness through modulation of lipid droplet. J Med Virol. 2023;95(3):e28630. doi:10.1002/jmv.28630

43. Liu T, Zhang L, Joo D, Sun SC. NF-κB signaling in inflammation. Signal Transduct Target Ther. 2017 Jul 14;2(1):17023. doi:10.1038/sigtrans.2017.23

44. Wang L, Cao JB, Xia BB, Li YJ, Zhang X, Mo GX, et al. Metatranscriptome of human lung microbial communities in a cohort of mechanically ventilated COVID-19 Omicron patients. Signal Transduct Target Ther. 2023 Nov 10;8(1):432. doi:10.1038/s41392-023-01684-1

45. Leon J, Michelson DA, Olejnik J, Chowdhary K, Oh HS, Hume AJ, et al. A virus-specific monocyte inflammatory phenotype is induced by SARS-CoV-2 at the immune–epithelial interface. Proc Natl Acad Sci. 2022 Jan 4;119(1):e2116853118. doi:10.1073/pnas.2116853118

46. Lian Q, Zhang K, Zhang Z, Duan F, Guo L, Luo W, et al. Differential effects of macrophage subtypes on SARS-CoV-2 infection in a human pluripotent stem cell-derived model. Nat Commun. 2022 Apr 19;13:2028. doi:10.1038/s41467-022-29731-5

47. Zhang J, Ejikemeuwa A, Gerzanich V, Nasr M, Tang Q, Simard JM, et al. Understanding the Role of SARS-CoV-2 ORF3a in Viral Pathogenesis and COVID-19. Front Microbiol. 2022 Mar 9;13:854567. doi:10.3389/fmicb.2022.854567

48. Apoorva, Shukla A, Kumar A, Singh S, Singh SK. Proteome profiling of human lung epithelial cells unveils distinct patterns of protein expression in response to SARS-CoV-2 ORF3a. The Microbe. 2025 Mar 1;6:100237. doi:10.1016/j.microb.2025.100237

49. Rui Y, Shen S, Wang Y, Cheng L, Chen S, Hu Y, et al. HIV-1 Vpu and SARS-CoV-2 ORF3a proteins disrupt STING-mediated activation of antiviral NF-κB signaling. Sci Signal. 2025 Jan 21;18(870):eadd6593. doi:10.1126/scisignal.add6593

50. Roquilly A, Jacqueline C, Davieau M, Mollé A, Sadek A, Fourgeux C, et al. Alveolar macrophages are epigenetically altered after inflammation, leading to long-term lung immunoparalysis. Nat Immunol. 2020 Jun;21(6):636–48. doi:10.1038/s41590-020-0673-x

51. Lercher A, Cheong JG, Bale MJ, Jiang C, Hoffmann HH, Ashbrook AW, et al. Antiviral innate immune memory in alveolar macrophages following SARS-CoV-2 infection ameliorates secondary influenza A virus disease. Immunity. 2024 Nov 12;57(11):2530-2546.e13. doi:10.1016/j.immuni.2024.08.018

52. Wang W, Qu Y, Wang X, Xiao MZX, Fu J, Chen L, et al. Genetic variety of ORF3a shapes SARS-CoV-2 fitness through modulation of lipid droplet. J Med Virol. 2023 Mar;95(3):e28630. doi:10.1002/jmv.28630

53. van Dierendonck XAMH, Vrieling F, Smeehuijzen L, Deng L, Boogaard JP, Croes CA, et al. Triglyceride breakdown from lipid droplets regulates the inflammatory response in macrophages. Proc Natl Acad Sci. 2022 Mar 22;119(12):e2114739119. doi:10.1073/pnas.2114739119

54. Singh A, Sen P. Lipid droplet: A functionally active organelle in monocyte to macrophage differentiation and its inflammatory properties. Biochim Biophys Acta Mol Cell Biol Lipids. 2021 Oct;1866(10):158981. doi:10.1016/j.bbalip.2021.158981

55. Lv S, Wang H, Li X. The Role of the Interplay Between Autophagy and NLRP3 Inflammasome in Metabolic Disorders. Front Cell Dev Biol. 2021 Mar 16;9:634118. doi:10.3389/fcell.2021.634118

56. Gao X, Zhu K, Qin B, Olieric V, Wang M, Cui S. Crystal structure of SARS-CoV-2 Orf9b in complex with human TOM70 suggests unusual virus-host interactions. Nat Commun. 2021 May 14;12:2843. doi:10.1038/s41467-021-23118-8

57. Jiang H wei, Zhang H nan, Meng Q feng, Xie J, Li Y, Chen H, et al. SARS-CoV-2 Orf9b suppresses type I interferon responses by targeting TOM70. Cell Mol Immunol. 2020 Sep;17(9):998–1000. doi:10.1038/s41423-020-0514-8

58. Thorne LG, Bouhaddou M, Reuschl AK, Zuliani-Alvarez L, Polacco B, Pelin A, et al. Evolution of enhanced innate immune evasion by SARS-CoV-2. Nature. 2022 Feb;602(7897):487–95. doi:10.1038/s41586-021-04352-y

59. Tan LY, Komarasamy TV, RMT Balasubramaniam V. Hyperinflammatory Immune Response and COVID-19: A Double Edged Sword. Front Immunol. 2021 Sep 30;12. doi:10.3389/fimmu.2021.742941

60. Lian Q, Zhang K, Zhang Z, Duan F, Guo L, Luo W, et al. Differential effects of macrophage subtypes on SARS-CoV-2 infection in a human pluripotent stem cell-derived model. Nat Commun. 2022 Apr 19;13:2028. doi:10.1038/s41467-022-29731-5

61. García-García T, Fernández-Rodríguez R, Redondo N, de Lucas-Rius A, Zaldívar-López S, López-Ayllón BD, et al. Impairment of antiviral immune response and disruption of cellular functions by SARS-CoV-2 ORF7a and ORF7b. iScience. 2022 Nov 18;25(11):105444. doi:10.1016/j.isci.2022.105444

62. López-Ayllón BD, de Lucas-Rius A, Mendoza-García L, García-García T, Fernández-Rodríguez R, Suárez-Cárdenas JM, et al. SARS-CoV-2 accessory proteins involvement in inflammatory and profibrotic processes through IL11 signaling. Front Immunol. 2023;14:1220306. doi:10.3389/fimmu.2023.1220306 PubMed PMID: 37545510

63. Love MI, Huber W, Anders S. Moderated estimation of fold change and dispersion for RNA-seq data with DESeq2. Genome Biol. 2014;15(12):550. doi:10.1186/s13059-014-0550-8

64. Ewels PA, Peltzer A, Fillinger S, Patel H, Alneberg J, Wilm A, et al. The nf-core framework for community-curated bioinformatics pipelines. Nat Biotechnol. 2020 Mar;38(3):276–8. doi:10.1038/s41587-020-0439-x

65. Di Tommaso P, Chatzou M, Floden EW, Barja PP, Palumbo E, Notredame C. Nextflow enables reproducible computational workflows. Nat Biotechnol. 2017 Apr;35(4):316–9. doi:10.1038/nbt.3820

66. Dobin A, Davis CA, Schlesinger F, Drenkow J, Zaleski C, Jha S, et al. STAR: ultrafast universal RNA-seq aligner. Bioinformatics. 2013 Jan 1;29(1):15–21. doi:10.1093/bioinformatics/bts635

67. Patro R, Duggal G, Love MI, Irizarry RA, Kingsford C. Salmon provides fast and bias-aware quantification of transcript expression. Nat Methods. 2017 Apr;14(4):417–9. doi:10.1038/nmeth.4197

68. Love MI, Soneson C, Hickey PF, Johnson LK, Pierce NT, Shepherd L, et al. Tximeta: Reference sequence checksums for provenance identification in RNA-seq. PLOS Comput Biol. 2020 Feb 25;16(2):e1007664. doi:10.1371/journal.pcbi.1007664

69. Galili T. dendextend: an R package for visualizing, adjusting and comparing trees of hierarchical clustering. Bioinformatics. 2015 Nov 15;31(22):3718–20. doi:10.1093/bioinformatics/btv428

