## Supplement figures for "Omicron-Enhanced Immunosuppressive Effects of SARS-CoV-2 ORF3a and ORF9b Accessory Proteins on Monocytic Inflammatory Response"

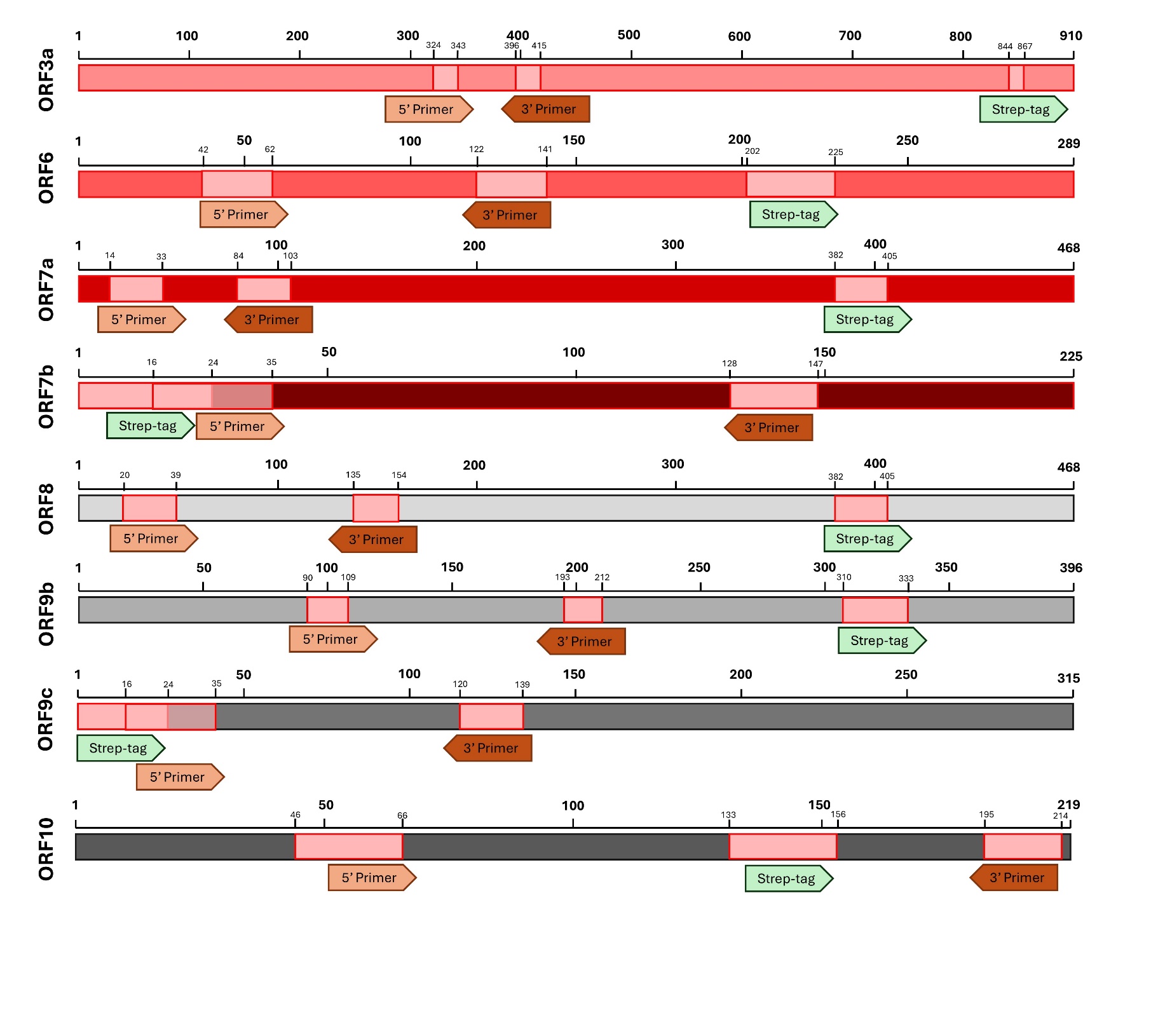

**Supplementary Fig. 1. SARS-CoV-2 accessory protein-coding sequences used for lentiviral vector construction and cell transduction.** Green arrows refer to 2x Strep-tag locations; brown arrows refer to forward and reverse primer locations for confirmation of insert presence with PCR (sequence length expressed in nucleotide base pairs (bp)).

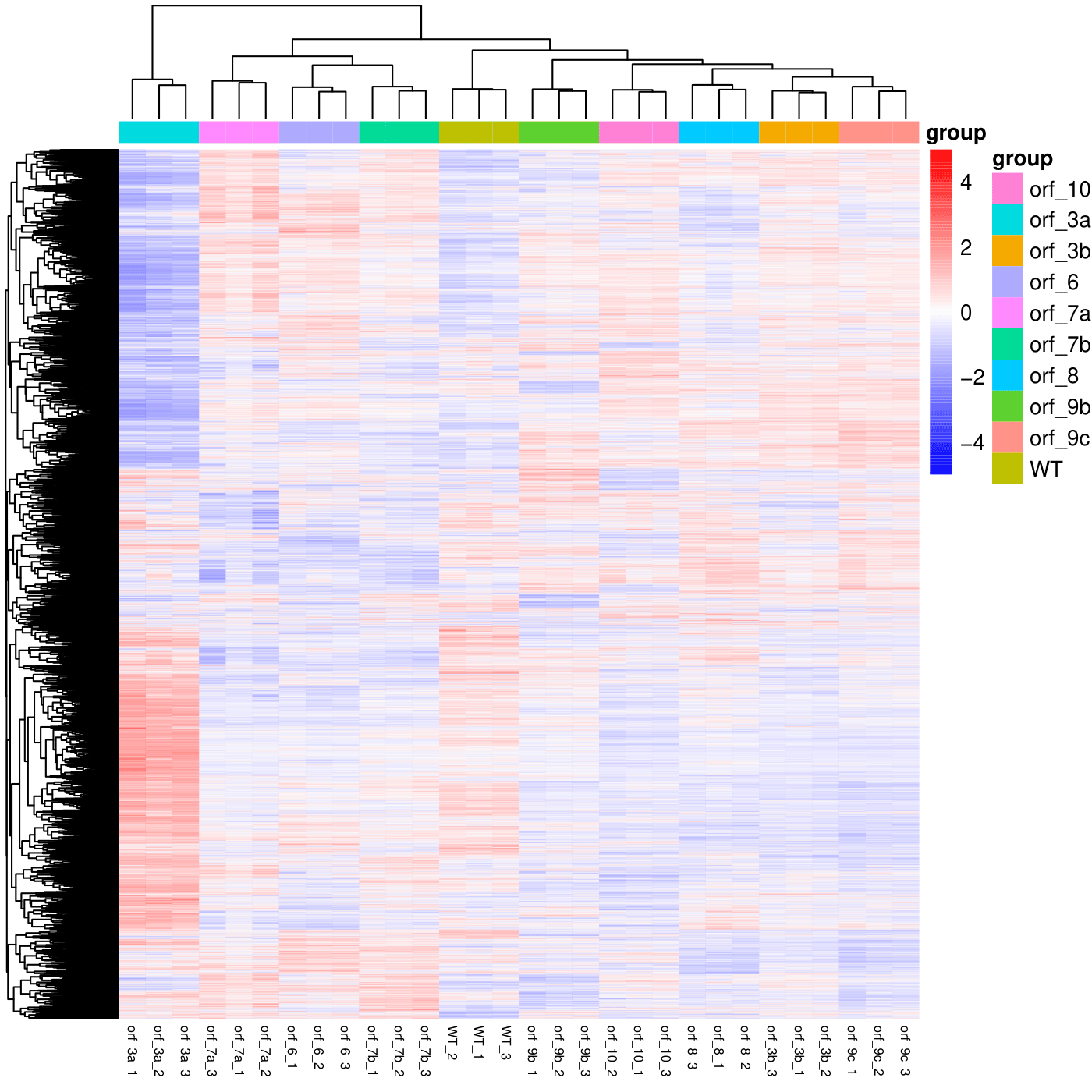

**Supplementary Fig. 2. A549-ORF cell transcriptome heatmap.** Gene count and clustering data presented for SARS-CoV-2 ORF3a-, ORF3b-, ORF6-, ORF7a-, ORF7b-, ORF8-, ORF9b-, ORF9c-, ORF10-transduced and control A549 (WT) cells in triplicates.

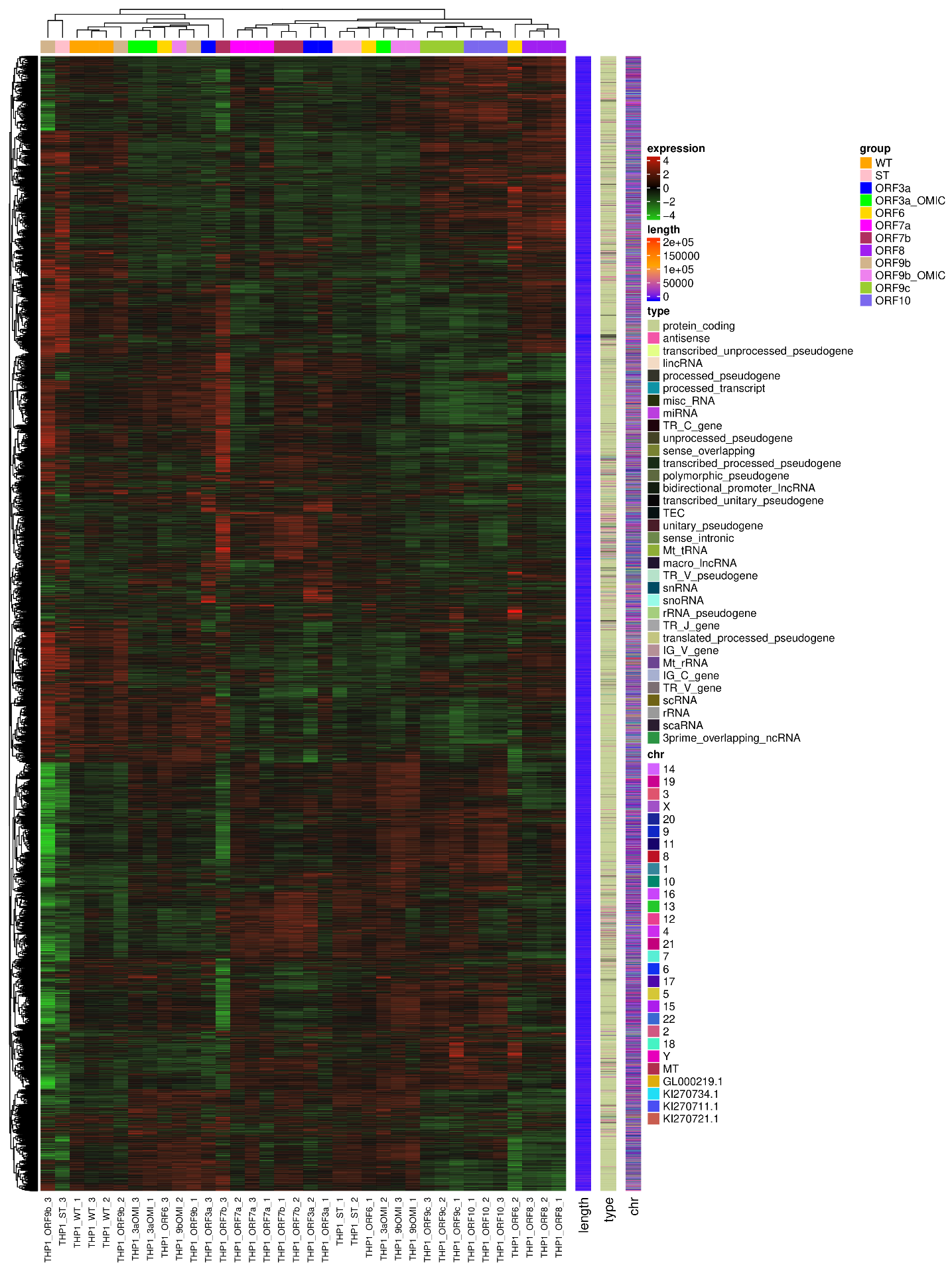

**Supplementary Fig. 3. THP-1-ORF cell transcriptome heatmap.** Gene count and clustering data presented for SARS-CoV-2 ORF3a- (Wuhan and Omicron origin), ORF6-, ORF7a-, ORF7b-, ORF8-, ORF9b- (Wuhan and Omicron origin), ORF9c-, ORF10-transduced and control A549 (WT and ST) cells in triplicates. Additionally, gene length, type and chromosome location data provided for each gene.

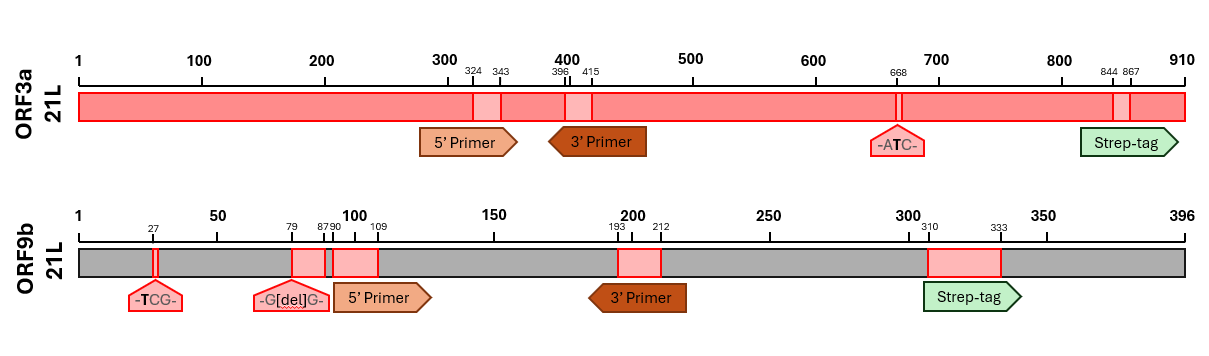
**Supplementary Fig. 4. SARS-CoV-2 BA.2 (21L) ORF3a and ORF9b-coding sequences used for lentiviral vector construction and cell transduction.** Green arrows refer to 2x Strep-tag locations; brown arrows refer to forward and reverse primer locations for confirmation of insert presence with PCR, red arrows indicate sites and types of mutations compared to Wuhan strains (letter in bold refer to point mutations; [del] refers to deletion) (sequence length expressed
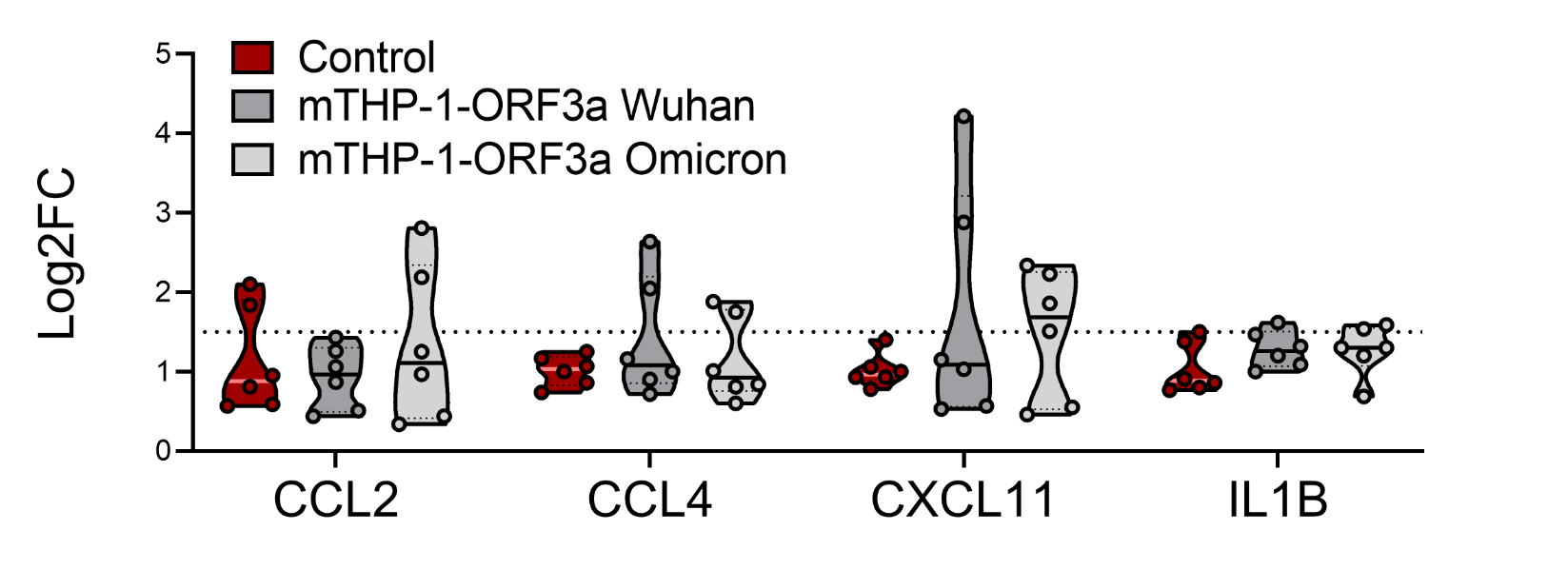

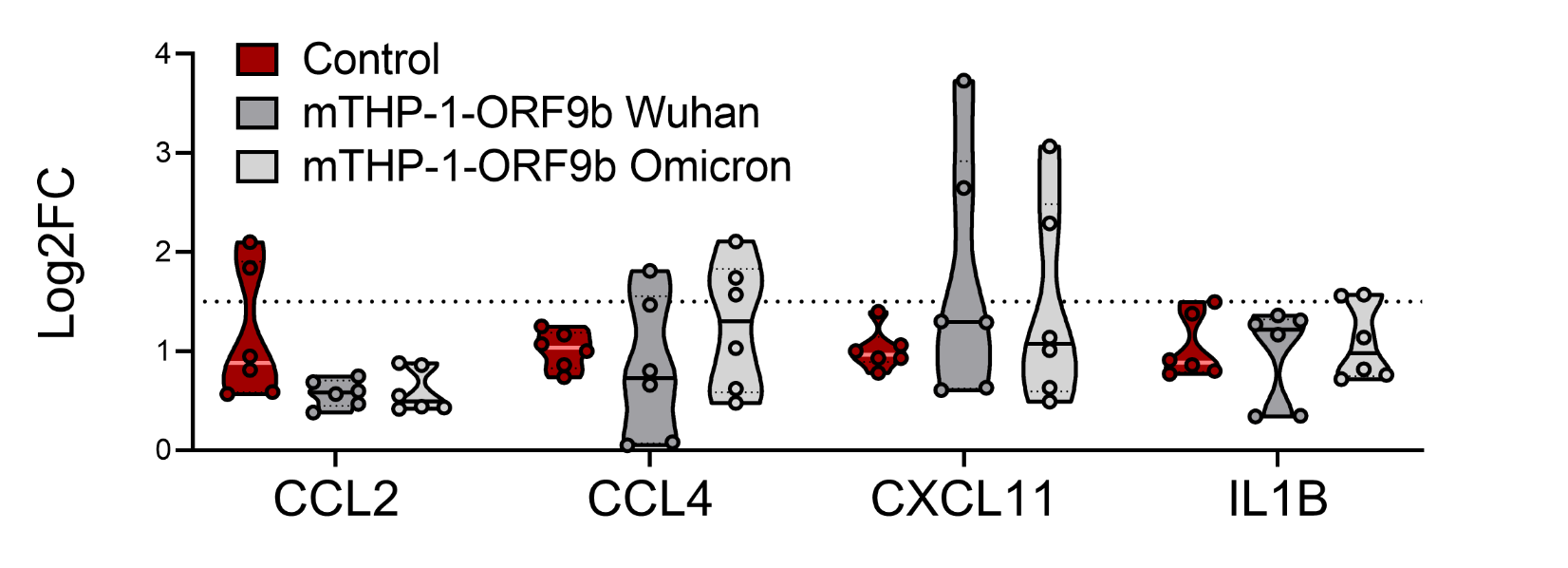
in nucleotide base pairs (bp)).

**A**

**B**

**Supplementary Fig. 5. Cytokine/chemokine mRNA expression in mock-stimulated control and SARS-CoV-2 ORF-transduced mTHP-1 cells.** Expression of *CCL2, CCL4, CXCL11* and *IL1B* mRNA in mock-stimulated Wuhan and Omicron **A** mTHP-1-ORF9b and **B** mTHP-1-ORF3a cells. Dotted line represents significantly altered expression (Log2FC > 1.5) relative to control mTHP-1 cell average. One-way ANOVA with Dunnett’s multiple comparisons test was used for comparison within each group.

| **Target** | **Forward** | **Reverse** |
| --- | --- | --- |
| GAPDH | CTGGTGGCTGGCTCAGAAAA | GGAGATTCAGTGTGGTGGGG |
| CCL2 | CCTTCATTCCCCAAGGGCTC | GGTTTGCTTGTCCAGGTGGT |
| CCL4 | GCTTCCTCGCAACTTTGTGG | TCACTGGGATCAGCACAGAC |
| CXCL11 | TTGCATAGGCCCTGGGGTAA | AGCCTTGCTTGCTTCGATTTG |
| IL1B | ACAGATGAAGTGCTCCTTCC | CGGCCTGCCTGAAGCCCTTG |
| ORF3a | GTACGCGCTCGTTTACTTCC | GAGGATTCTTGCTCCGACAC |
| ORF3d | GCATACTGCTGGAGATGCAC | GCAGGCTATTGCATACAACG |
| ORF6 | ACTCCTCATAATCATGCGGAC | ATTCTCGGTCAGGCTCTTGG |
| ORF7a | TGTTTCTGGCCCTCATAACC | AGGGCTCTTTCAGCAGTACG |
| ORF7b | TTTGAGAAGGGTGGTGGTTC | AAGAAGGAAGGCGAGGAAAC |
| ORF8 | TGGGCATTATCACCACAGTC | GGGCCCCTACCCTGATATAC |
| ORF9b | AGGACGCGACCAAAATAATG | AGCTGAAACGCTTTGTCCTC |
| ORF9c | TTTGAGAAGGGTGGTGGTTC | TTTGGCAGTGTTGCTCTTTC |
| ORF10 | CTCTTGTTGTGTCGCATGAAC | TTTCAAACTGCGGATGTGAC |
| Control 2x Strep-tag | TTTGAGAAGGGTGGTGGTTC | GCAGCGCCTTTTTCAAATTGC |

**Supplementary Table 1. Primer pairs used for qRT-PCR**. Targets include housekeeping gene GAPDH, cytokine/chemokine mRNA (CCL2, CCL4, CXCL11 and IL1B), SARS-CoV-2 accessory proteins (ORF3a, ORF3d, ORF6, ORF7a, ORF7b, ORF8, ORF9b, ORF9c and ORF10) and 2x Strep-tag for transduction evaluation.

**Supplementary information**

Supplementary information for: Omicron-Enhanced Immunosuppressive Effects of SARS-CoV-2 ORF3a and ORF9b Accessory Proteins on Monocytic Inflammatory Response

The file includes Supplementary Figures 1-5 and Supplementary Table 1.

**Supplementary Fig. 1.** SARS-CoV-2 accessory protein-coding sequences used for lentiviral vector construction and cell transduction.

**Supplementary Fig. 2.** A549-ORF cell transcriptome heatmap.

**Supplementary Fig. 3.** THP-1-ORF cell transcriptome heatmap.

**Supplementary Fig. 4.** SARS-CoV-2 BA.2 (21L) ORF3a and ORF9b-coding sequences used for lentiviral vector construction and cell transduction.

**Supplementary Fig. 5.** Cytokine/chemokine mRNA expression in mock-stimulated control and SARS-CoV-2 ORF-transduced mTHP-1 cells.

**Supplementary Table 1.** Primer pairs used for qRT-PCR.
